# Chemotactic interactions drive migration of membraneless active droplets

**DOI:** 10.1101/2023.04.25.538216

**Authors:** Mirco Dindo, Alessandro Bevilacqua, Giovanni Soligo, Alessandro Monti, Marco Edoardo Rosti, Paola Laurino

## Abstract

In nature, chemotactic interactions are ubiquitous and play a critical role in driving the collective behaviour of living organisms. Reproducing these interactions *in vitro*is still a paramount challenge due to the complexity of mimicking and controlling cellular features, such as metabolic density, cytosolic macromolecular crowding and cellular migration, on a microorganism size scale. Here we generate enzymatically-active cell-size droplets able to move freely and, by following a chemical gradient, able to interact with the surrounding droplets in a collective manner. The enzyme within the droplets generates a pH gradient that extends outside the edge of the droplets. We discovered that the external pH gradient triggers droplet migration and controls its directionality, which is selectively towards the neighbouring droplets. Hence, by changing the enzyme activity inside the droplet we tuned the droplet migration speed. Further, we showed that these cellular-like features can facilitate the reconstitution of a simple and linear protometabolic pathway with improved overall activity. Our work suggests that simple and stable membraneless droplets can be applied to reproduce complex biological phenomena opening new perspectives as bioinspired materials and synthetic biology tools.

## Introduction

Cellular migration represents a key feature of living systems [1–3]. It manifests as the response to a stimulus which drives the directionality of the cellular motility. The migration can be mediated by chemotactic sensing, which plays a critical role in many processes, driving long-range interactions and collective behaviors of living systems, as well as sustaining and regulating life [1]. Chemotactic strategies have been reported to drive cell migration towards a higher chemoattractant molecule concentration, such as glucose, amino acids and other cell nutrients (positive chemotaxis) [4], or towards a lower chemorepellent molecule concentration, such as phenol and other cytotoxic compounds (negative chemotaxis) [5]. Chemotactic interactions also play a significant role in multicellular life, e.g. they are involved in organism development [6], immune response [7], cancer [8, 9] and inflammatory diseases [10]. Due to its fundamental role, current intracellular pathways leading to cellular chemotaxis have been deeply studied [11, 12], but mimicking *in vitro* the cellular degree of complexity is challenging. In fact, how nature has achieved this level of complexity is not completely understood [13, 14]. Studying how these cellular functions work and arise require life-like systems capable of mimicking cellular complex features, such as metabolic density, cytosolic macromolecular crowding, proliferation and migration [15–17]. Thus, reproducing pivotal chemotactic interactions is a key step to unravel basic natural rules driving collective behavior and the physical principles ruling these phenomena.

The design and control of synthetic droplets able to recall cellular features and respond to chemical signals represent a new frontier for intelligent materials [18–20] and biotechnology [21–27]. In general, synthetic droplets can be obtained either by liquid-liquid phase separation or by encapsulating their content in a membrane (vesicles) [28–38]. Synthetic droplets have been deeply exploited to study cellular features [39–41] or to reproduce them [18, 42–44]. These studies highlight the potential of using synthetic droplets systems to study biologically relevant questions, but also the current limitations and challenges in mimicking the complexity of living systems [45]. In detail, the presence of a surrounding membrane strongly affects, or impedes completely, the diffusion of molecules in and out of the vesicles, limiting studies on chemotactical sensing [46] or enzyme kinetics [47]. On the other hand, membraneless liquid droplets are affected by external environmental factors, such as pH, ionic strength or temperature [48]. For these reasons, they show poor stability (dissolution, spreading on the surfaces and aggregation) and they do not usually respond to chemical gradients [49]. In conclusion, these studies do not report droplets able to respond to a chemical signal by showing migration triggered by chemotactic sensing.

In this work, we report how enzymatically-active droplets respond and move when triggered by a self-produced chemical gradient of pH in a chemotactic manner. The droplet microenvironment not only mimics intracellular features like cytosolic protein crowding, metabolic activity and cell-like length scale (Fig. 1A), but also resembles intercellular complex interactions, such as migration and merging. The enzyme partitioned in the droplets generates a pH gradient which extends outside the droplets and modifies the surface tension of the droplets according to the local pH. Changes in the surface tension value along the interface of the droplets generate stresses tangential to the interface, known as Marangoni stresses. The Marangoni-induced flow is able to propel the droplet towards the direction that minimizes its surface energy [50], i.e. towards higher values of pH, represented by the neighbouring active droplets (Fig. 1B). In summary, droplets influence, sense each other and move towards the neighboring ones driven by a difference in pH generated by the enzymatic activity. We performed numerical simulations to corroborate the experimental observations, confirming the dynamics of the motion of the droplets. When the self-generated chemical gradient is modified, by varying substrate and enzyme concentrations, it tunes the droplet migration speed; making this system the first proof-of-concept for the design of controllable moving system based on enzymatic activity (Fig. 1C). Interestingly, we also demonstrate that our system can trigger and facilitate the formation of a simple linear metabolic pathway, providing a possible mechanism for long-range interactions in droplets communities.

**Fig. 1.**
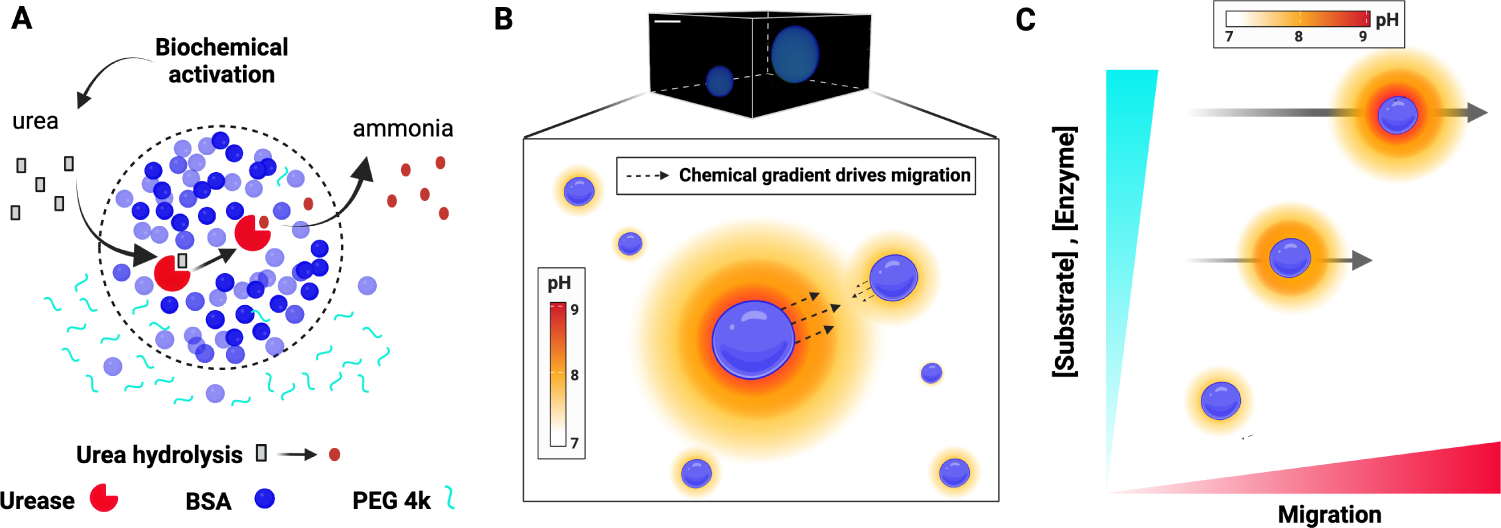
Active membraneless droplets show migration governed by an enzyme-generated chemical gradient. **A** Scheme of the liquid-phase separated system employed in the paper. Active droplets containing urease efficiently catalyzed the conversion of urea to ammonia. **B** Up to down. Z-stack visualization of the droplets containing Alexa Fluor 488 labeled-BSA (colored in blue) using PEGDA 10 kDa as a glass slide coating and schematic representation of pH driven droplets migration (right). Active urease-containing droplets (1-1.5 *µ*M) after adding 115 mM urea are able to generate a pH gradient which affects the droplet surface tension accordingly and ultimately inducing migration. Scale bar 20 *µ*m. **C** Scheme of the modulation of the droplets migration obtained by varying substrate and enzyme concentration.

## Results

### Generating non-adhering droplets on a glass surface

To avoid glass-adhering phenomena and dissolution of the droplets on the glass slide, several polyethylene glycol diacrylate polymers (PEGDA) were screened as glass slide coatings. The tested polymers were PEGDA 700 Da, PEGDA 4 kDa, PEGDA 6 kDa and PEGDA 10 kDa [51]. In details, we imaged Alexa Fluor 488-labeled Bovine Serum Albumin (BSA) droplets using a confocal microscope and we calculated the contact angle formed between the droplet and the coated glass by analyzing 3D images of sessile droplets using the contact angle plug-in of ImageJ software [52]. This analysis measured a contact angle *θ* (theta) which increases from 114*^◦^ ±* 5*^◦^* in PEGDA 700 Da to 149*^◦^±*6*^◦^* using PEGDA 10 kDa (Fig. S1). PEGDA 10 kDa showed the least adhesion of the droplets on the glass slide coating, resulting in spherical droplets that show no resting phenomena on the PEGDA-glass slide surface. For these reasons, PEGDA 10 kDa was chosen as glass slide coating for this study.

### Enzymatic activity generates and control the pH gradient around the droplets

Enzymatically-active urease-containing droplets (1-1.5 *µ*M) (partitioning is reported in Fig. S2) [53] are able to generate a pH gradient inside the droplets and in the nearby external vicinity of the droplet’s interface, that we will call herein as the droplets’ halo. The basic pH gradient inside and outside the droplets is due to the production of ammonia as a product of urea hydrolysis catalyzed by urease (Fig. S3). We previously studied the droplets inner pH gradient [53]. Here, we focus on the study of the pH halo around the droplets, and on its impact on the droplets and surrounding solution. By adding 50-100 *µ*M of the pH sensitive probe SNARF-1 in the supernatant phase along with a saturating concentration of urea (115 mM), we measured and quantitatively analyzed the halo features (as intensity of the signal or as pH values) of the reacting droplets. As reported by the raw images in Fig. 2A and by the quantitative analysis (Fig. 2B), the fluorescence emission intensity of SNARF-1 as well as the pH values around the enzymatically active droplets are strictly related to their overall size. Indeed, the active droplets with radius of 40-50 *µ*m shows higher pH values compared to those of the smaller droplets (radius of 8-15 *µ*m). Further, we have analyzed the pH amplitude of the halo around the reacting droplets of different size (Fig. 2C). Large active droplets (radius of 60 *µ*m) show high internal pH values (Fig. S4) which propagate around their edges on the surrounding environment for several micrometers (*>*100 *µ*m). Smaller active droplets (radius 8-50 *µ*m) instead show negligible differences between the inner and outer pH (Fig. 2C). In addition, in agreement with the data reported in Fig. S4, the extension of the pH halo is strictly correlated to the droplet’s size. Despite the intensity of the halo signal decreases by increasing the distance from the droplet edges (Fig. 2D), it remains stable for several minutes (Fig. 2E). Lastly, we analyzed the behavior of two different active droplets (Fig. 2F). The single droplet analysis shows that the small droplet (r = 22 *µ*m) does not migrate. In fact, it shows a modest internal pH change (Δ pH*≤*0.5) which is accompanied by a negligible pH halo around its edges. On the other hand, the large droplet (r = 64 *µ*m) does show migration. Interestingly, this droplet shows a high internal pH change (Δ pH*≈*2) which generates a wide (*>*50*µ*m) pH halo around its edges. These data indicate that the pH halo intensity and amplitude is strongly related to the size and the pH change of the droplet, and that it can stably propagate itself into the surrounding environment for several micrometers. This key feature allows the droplets to “sense” through chemotactic interactions the nearby droplets, driving the directional migration.

**Fig. 2.**
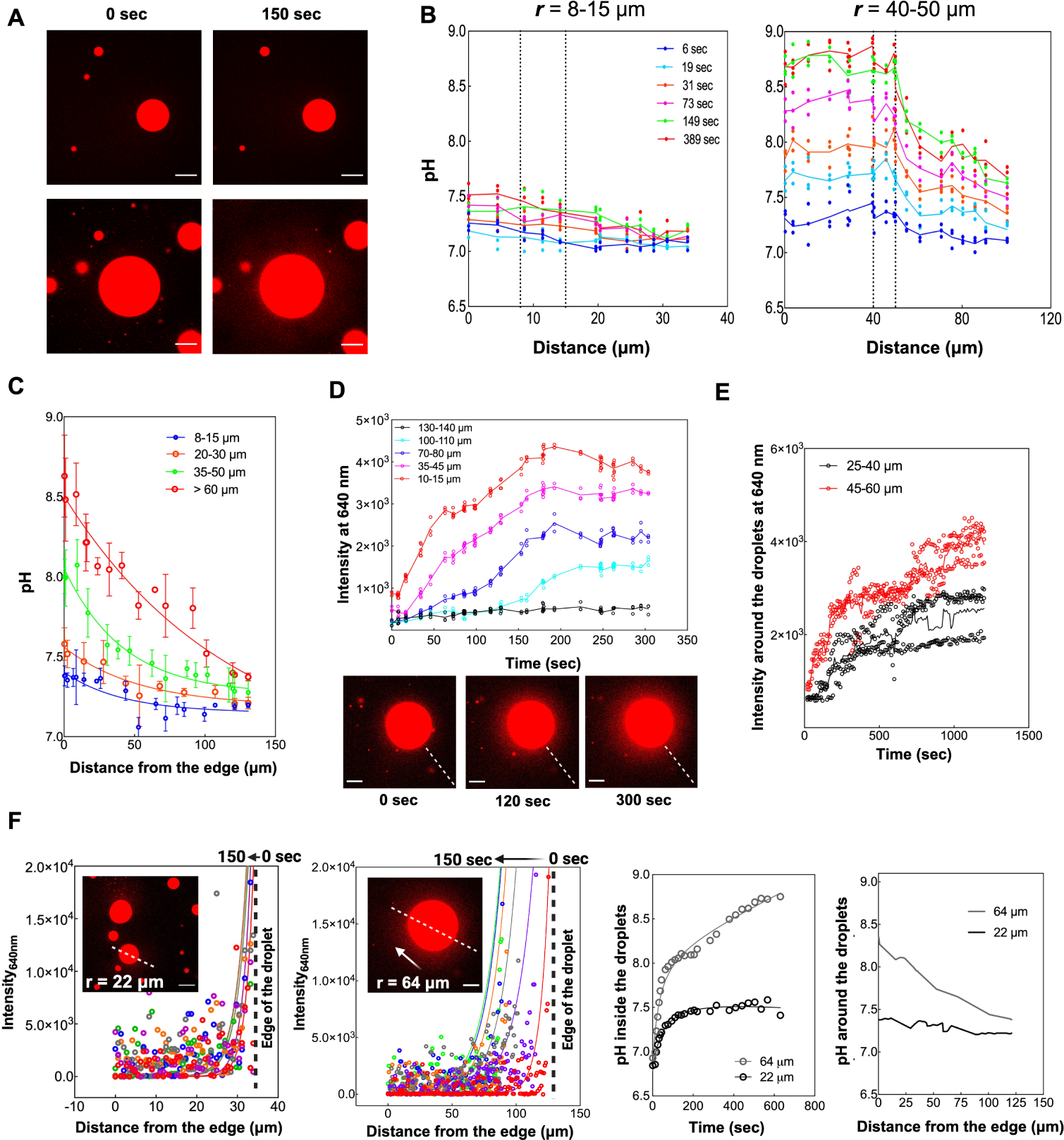
Analysis of the halo around the active droplets reveals its features and highlight its importance for driving migration. **A.** Raw images of pH-sensitive probe SNARF-1 used both labeled to BSA and free in the supernatant phase (50-100 *µ*M) for the visualization and the analysis of the internal and external pH changes of the reacting droplets at different times. Droplets in the images contain 1-1.5 *µ*M urease upon addition of 115 mM urea. Individual color channel (red) adjustment have been performed to highlight the pH change. **B.** Quantitative evaluation of the pH halo change over distance from the droplet center (*µ*m) at different times for droplets with different radius (around 8-15 *µ*m and around 40-50 *µ*m). Free SNARF-1 in the supernatant phase were used to determine the pH outside the (pinned) reacting droplets by using the calibration curve reported in Fig. S4. Dotted vertical lines represent the droplets edges. Data are represented with at least three different droplets in the size range reported (mean values *±*SD) **C.** Quantitative pH halo analysis over distance around pinned reacting droplets, clustered based on their different radius. The measurements have been done after 200-250 seconds starting from the edges of the reacting droplets over a distance of 135 *µ*m. Droplets analyzed contain 1-1.5 *µ*M urease upon addition of 115 mM urea. **D.** Trend of free SNARF-1 emission fluorescence intensity at 640 nm over time at different distance from the edge of pinned reacting droplets (droplets radius analyzed is between 45-62 *µ*m, urease concentration 1.0 *µ*M, urea concentration 115 mM). Raw microscope qualititive images of the halo progression over time of a single droplet (r = 62 *µ*m). **E.** Measurement of halo stability over time. Analysis of SNARF-1 emission fluorescence intensity at 640 nm around pinned reacting droplets (measured as intensity of free SNARF-1 in the supernatant) of different radius (clustered as r = 25-40 *µ*m and 45-60 *µ*m). The intensity of the signal around the droplets (distance was 10-15 *µ*m from the edge) is stable for more than 10-20 minutes after the pH inside the droplets have reached the plateau. **F.** Qualitative measurement of the pH halo of an active moving and active not moving droplet. Left to right. Fluorescence emission intensity at 640 nm of free SNARF-1 for a droplet (r = 22 *µ*m) which does not display migration. Inset: raw image of the analyzed selected droplet. Fluorescence emission intensity at 640 nm of SNARF-1 for a droplet (r = 64 *µ*m) which shows migration, by following the direction indicated by the white arrow. The dotted black line on the graphs indicates the edge of the droplets. Internal and external pH profiles of the two reacting droplets analyzed. All the experiments shown have been performed using an overall concentration of urease in the droplets of 1.0 *µ*M and substrate urea equal to 115 mM. All the experiments have been performed at least in triplicate except for the qualitative measurements of the halo in the moving droplets (panel F). The imaging experiments using SNARF-1 have been performed by using a Zeiss Airscan LSM-880. Scale bars are 50 *µ*m.

### Migration is driven by a chemical gradient

The pH change generated via enzymatic activity drives the motion of the droplet: the change of pH determines a change in surface tension [53], which in turn induces Marangoni stresses. These stresses act tangentially to the interface of the droplets and are directed from low-surface tension regions towards high-surface tension regions, and generate a re-circulatory motion inside the droplets [53], which then propels the droplet itself in the direction of its low-surface tension pole, minimizing its total energy [50]. Flow visualization using fluorescent nanoparticles showed the Marangoni-induced flow inside the droplets (Supplementary movies 1-2).

A change of pH is fundamental in the migration of the droplets: we confirmed that no motion is recorded in the absence of enzymatic activity generated by urease and its substrate, urea. The absence of the substrate urea in the urease-containing droplets (1-1.5 *µ*M), generates no migration (Supplementary Movie 3) and no flow inside the droplets [53]. Without urease in the droplets, no migration was detected after adding 115 mM urea in the droplet solution (Supplementary Movie 4). Furthermore, if other enzymes that do not produce any detectable pH change were partitioned in the droplets, as lactate dehydrogenase and glucose oxidase (reaction scheme in Fig. S3 and partitioning Fig. S2), no motion of the droplets was observed (Supplementary Movie 5 and 6).

We have thus identified Marangoni stresses as the driving mechanism of the migration of the droplets. For Marangoni stresses to be present two conditions must be verified: (*i*) a pH gradient along the interface of the droplet must be present and (*ii*) the pH change must correspond to a (significant) surface tension change. We will first discuss how a pH gradient along the interface can be formed. The chemical reactions occurring inside the droplet generate a pH halo surrounding the droplet itself, as shown in Fig. 2. A single, isolated droplet produces a uniform halo all around the droplet itself and the pH value at the interface is uniform. To achieve a pH gradient along the interface we must introduce a perturbation in the pH field: the interaction with the halo generated by another nearby droplet for instance generates a non-uniform pH value at the interface of the droplet. It is challenging to verify this condition in experiments as droplets in the far field introduce perturbations in the pH field; it is instead easily achieved in numerical simulations, where we observe that single, isolated droplets do not exhibit any motion due to the absence of Marangoni stresses, as the pH value is uniform along the interface. This can be appreciated in our numerical simulations, Fig. 3 panels B and C, where upon coalescence of the droplets, the single merged droplet stops moving. Conversely, in Fig. 3D, upon the first merging, the newly-formed droplet keeps moving: the interaction with the halo generated by the surrounding droplets causes an inhomogeneous pH value (thus, an inhomogeneous surface tension) along the interface, inducing Marangoni stresses along the interface of the droplets, and consequently driving the droplets towards each other. The pH gradient along the interface thus attained then must induce a change in surface tension large enough to generate Marangoni stresses at the interface. The change of surface tension for varying values of pH was measured in a previous work [53] and their experimental measurements are reported in the inset of Fig. S8, where a strong reduction in surface tension for values of pH larger than pH*>* 7.5 is observed. Upon addition of a saturating concentration of the substrate urea (115 mM), urease-containing droplets with radius *<* 25 *−* 35 *µ*m reach an internal pH value of about 7.5 at most, and thus show no migration (Fig. 3A and Fig. S7). The change of surface tension in this range of pH is too low to produce significant Marangoni stresses, see the inset of Fig. S8, and thus no motion is observed. To further test this finding, another enzyme, cystalysin (reaction scheme Fig. S3 and partitioning Fig. S2), was used. A similar pH change was observed in the range pH=[6.5 : 7]; again this change of pH does not produce any significant change in the surface tension, even for droplets larger than *>* 70 *µ*m (Fig. S5). Another aspect to consider is linked to the strength of the halo: the pH inside the droplet depends on its size, see Fig. S4. Larger droplets have a higher internal pH value and thus the halo extends further away from the interface of the droplet Fig. 2C. When considering smaller droplets, such as those in Fig. S7, we do not observe any motion: the pH inside these (in the range of radius *≈*10-40 *µ*m) is rather low (pH*≈*7 -7.5), thus the width of the halo is limited and, as already discussed earlier, the change in surface tension along the interface is minimal. Marangoni stresses, in this case, are too weak to produce coherent motion of the droplet.

**Fig. 3.**
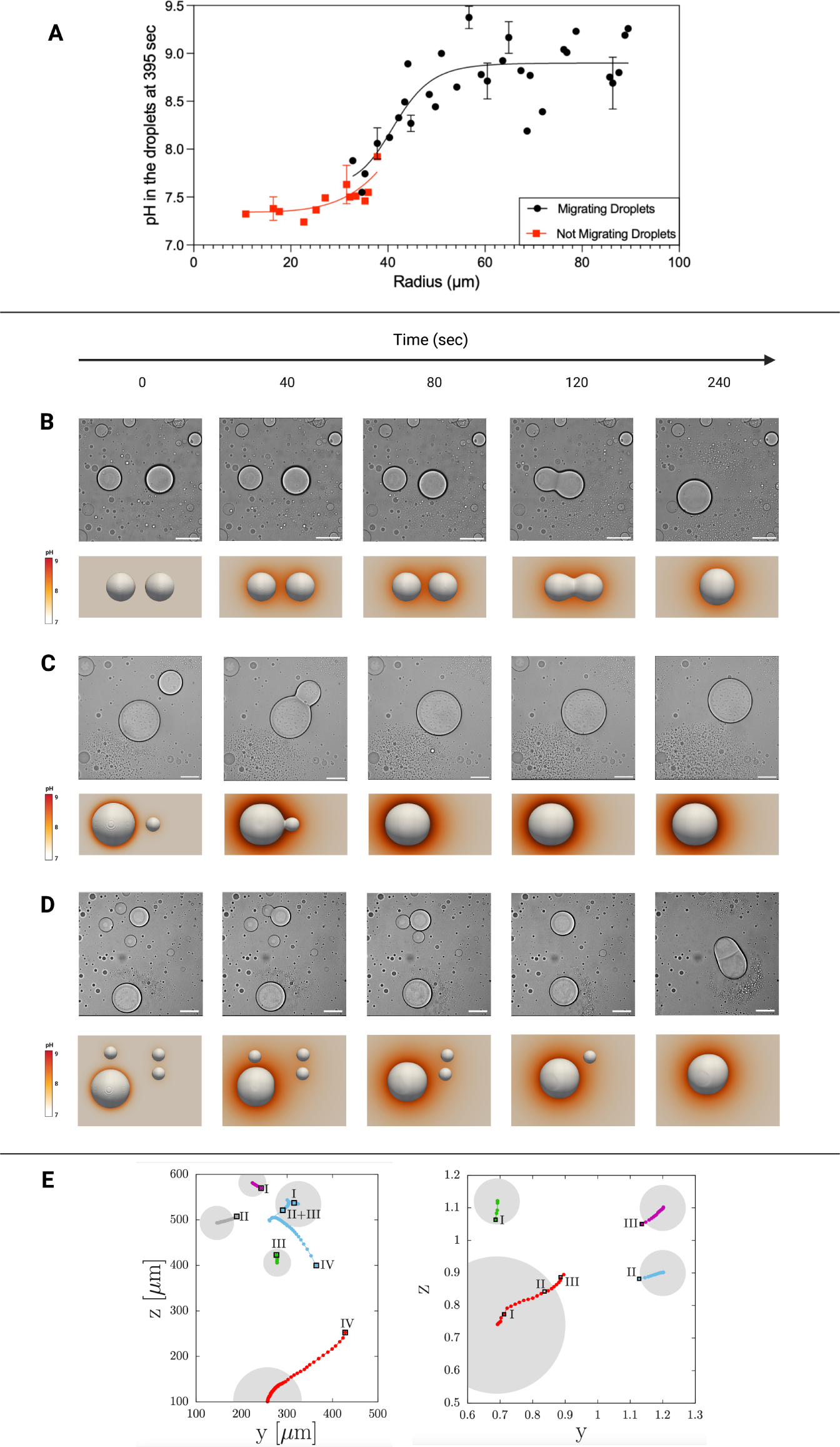
The migration mediated by the pH change of the active droplets is influenced by the droplet’s size. **A.** The graph shows the relation between droplets size and their internal pH. Red squares represent the droplets which do not show migration, while black dots highlight the droplets which show migration. Lines represent the sigmoidal fitting of the two datasets. More than 50 active droplets with different size were analyzed which contain 1-1.5 *µ*M urease in presence of the substrate urea (115 mM). Data from droplets of the same size are averaged (mean value *±*SD). **B-D.** Top: visualization at different times of similar size (B), of different size (C) and of multiple (D) active droplets containing 1-1.5 *µ*M urease in presence of 115 mM urea which show migration and merging in solution. The snapshots have been obtained by analyzing the movies present in the Supporting information (Supplementary Movie 7, 8 and 9) at different times. Scale bar: 100 *µ*m. Bottom: numerical simulations of the droplets supporting the experimental evidence (Supplementary Movie 12, 13 and 14); the simulation snapshots shown are taken at different times. **E.** Trajectory analysis performed by tracking the center of the droplets of the experiment (left) and numerical simulation (right) reported in panel D. Different colors represent the center of different droplets over time; markers are drawn every Δ*T* = 6 sec (experiment, left) and Δ*T* = 0.04 (simulation, right). The droplets’ trajectory have been analysed using MATLAB software. Each merging event is numbered (I-IV, experiment and I-III simulation) and highlighted with black empty squares.

These tests confirm that Marangoni stresses are the mechanism driving droplet’s motion, and no motion is observed in the absence of Marangoni stresses. The droplet motion ultimately originates from the presence of pH gradients at the interface of the droplet itself and can be viewed as a form of chemotaxis.

In Fig. 3B-D we report several snapshots from our experiments and numerical simulations representative of three sample cases: (i) two droplets of (approximately) the same size (panel B), (ii) two droplets of different sizes (panel C) and (iii) several droplets of different sizes (panel D). In the experiments it is very challenging to obtain binary systems of droplets, in which there is no long-range effect from other droplets further away; numerical simulations allow us to single out these far field effects and focus solely on droplet-droplet interactions. Thus, We will use data from numerical simulations to show the evolution and interaction of the halos surrounding the droplets, which proved to be fundamental in determining the motion of the droplets.

In Fig. 3B and Supplementary Movie 7 we analyse the case where two droplets of approximately the same size (50 *µ*m radius in the experiment) are placed close to each other (initial distance of 90 *µ*m in the experiment). Snapshots from the numerical simulations show the evolution of the pH halo around the droplet; the pH halo starts diffusing outwards from the droplets (up to roughly 40 sec, experiment time). Due to the interaction between the halos of the two droplets, the pH is higher in between them, thus generating an imbalance in the pH value along the interface of the droplets. The surface tension gradient due to the pH imbalance generates Marangoni stresses, and ultimately drives the droplets towards each other. The experimental snapshots show that these droplets merge over a *≈* 120 sec time-scale. The lower pH value inside the droplets generates a relatively weaker halo (i.e. extending less off the interface and generating smaller perturbations at the interface of other droplets) compared to larger droplets.

The effect of the droplet size can be understood from the cases shown in Fig. 3C and Supplementary Movie 8, with two droplets of different sizes (radius equal to *∼*110 *µ*m and *∼*60 *µ*m, respectively) placed at a relatively close distance, similar to panel B, *≈*100 *µ*m. Similar dynamics are observed for this case as well: the halo diffuses outwards from the droplets in approximately the initial 40 sec (experiment time) and a higher value of pH is found in between the droplets (right pole of the large droplet and left pole of the small droplet). The pH, and thus surface tension, inhomogeneity generates Marangoni-induced flowing which propels the droplets towards each other. The merging occurs within a much shorter time interval, about 40 sec, compared to the case reported in Fig. 3B (about 120 sec). Here the pH halo generated by the larger droplet is wider and higher (according to Fig. 2), thus speeding up the overall dynamics, as shown in Fig. S10. We observe that also the smaller droplet moves: while the internal pH of the droplet is too low to generate motion (i.e. pH*≤* 7.5), the pH halo from the larger droplet causes a pH hotspot at the right pole, with local pH values larger than pH= 7.5. Upon merging, the droplet in the experiments slowly drifts away, due to the long-range effect of other, out-of-field droplets. In the numerical simulations we do not have effects from other droplets: upon merging there is no longer any imbalance in the halo around the droplet, thus no Marangoni stresses are present. The newly formed droplet does not move any further.

In the third sequence in Fig. 3D and Supplementary Movie 9, we study a multiple-drop case: several droplets of different sizes (and thus different internal pH values) interact with each other. We observe both in our experiments and numerical simulations that larger droplets (radius ≥ 50 *µ*m) are those that cover the longest distance and merge into the smaller droplets (radius *≈* 30 *µ*m). The halos generated by the larger droplets extends further away from their interface, and perturbs the pH concentration at the interface of the nearby droplets; closer droplets are clearly subjected to stronger perturbations. The inhomogeneous pH distribution along the interface of the droplets generates a Marangoni-induced flow, which propels the droplets toward each other. When the droplets are close to each other, the pH perturbations at the interface of both large and smaller droplets (here the threshold is about 25-35 *µ*m in radius) is able to generate a non-homogeneous surface tension along the interface, and thus Marangoni-driven self-propulsion. Conversely, when the droplets are further apart, the halos from other surrounding droplets can modify the surface tension (and thus inducing Marangoni-driven motion) only in the larger droplets, i.e. radius ≥ 50 *µ*m. This is also shown in the simulation snapshots: the two smaller droplets on the right are relatively close to each other but will not move until the larger droplet is close enough. This results further corroborates the observation that smaller droplets do not exhibit migration, unless they are in the near proximity of larger ones, as reported in Fig. S7. The larger droplets instead propel themselves according to the surrounding pH gradient, which is given by the superposition of the halos from all the surrounding droplets. These qualitative observations are backed by the trajectories of the center of the droplets, Fig. 3E. Markers on the trajectories are plotted at regular time intervals, thus providing a visual quantification of the speed of the droplets. In both the experiment and numerical simulations, the larger droplets are those covering the longer distances, moving at first towards the closest droplet(s), merging with them and then continuing in their motion for as long as there is an imbalance in the surface tension along their interface. We observe that smaller droplets start moving only when in the near proximity of larger ones, as the latter ones generate a perturbation in the pH field strong enough to significantly modify the surface tension at the interface.

### Substrate and enzyme concentration tune droplet migration

Here, we show how the droplets migration displayed in the previous section is tuned by the enzyme activity of the droplets . Specifically, migration speed is controlled by changing either enzyme or substrate concentrations (Fig. 4). In Fig. 4A and Fig. S6, we have calculated the trajectories of the active droplets over time using several substrate concentration (0-200 mM). The analysis of the trajectories represents the distance travelled by the droplets over time (r *>* 35 *µ*m) when subjected to different substrate concentrations (Fig. 4B). In detail, the addiction of 0 to 5 mM urea results in no droplet migration. By increasing the substrate concentration (from 10 to 200 mM) the droplets gradually start to migrate, showing a linear trend between the distance travelled and the substrate concentration over time. These data clearly show that the modulation of the substrate concentration regulates the migration features of the active droplets. Interestingly, migration control can be achieved by changing the enzyme concentration. For instance, by increasing the urease concentration within the droplets from 1 *µ*M to 3.3 *µ*M (Fig. 4C). Using a fixed urea concentration 115 mM), enzymatically active droplets containing 3.3 *µ*M cover longer distances compared to droplets containing 1 *µ*M. Lastly, a quantitative analysis between distance covered by active droplets and their size at different substrate concentration is reported in Fig. 4D. The trajectories show that the migrated distance increase accordingly with the droplet size; even though for droplets of radius *>* 50 *µ*m this effect slightly decreases, probably because both the internal and external pH have reached the plateau (Fig. S4).

**Fig. 4.**
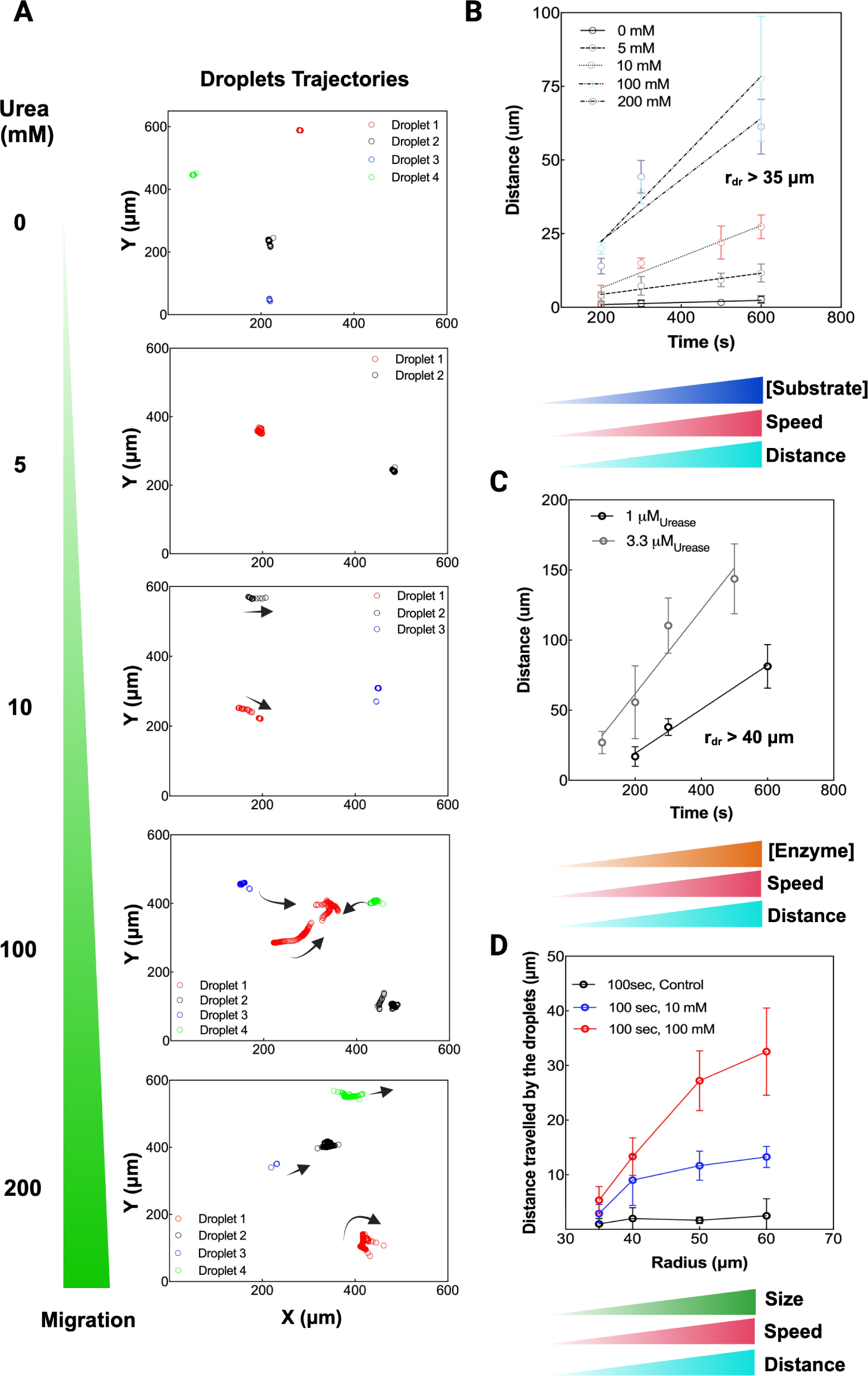
Droplets migration is tuned by varying substrate and enzyme concentrations. **A.** Top to bottom: droplet trajectory analysis performed by tracking the center of each droplet obtained by varying substrate concentration from 0 to 200 mM urea, while enzyme (urease) concentration was 1 *µ*M. The droplets’ trajectory has been analyzed using MATLAB software and every marker is drawn every Δ*T* = 2 sec. The direction of the movement is indicated in all the images by the arrows. All the droplets analyzed have a radius *≥* 35 *µ*m. No merging events are highlighted in this panel. More trajectory images are reported in Fig. S6. **B.** Linear correlation between the distance traveled by the droplets over time at different substrate concentrations. **C.** Linear correlation between the distance traveled by the droplets over time obtained at two different enzyme concentrations. Trajectory analysis is reported in Fig. S6. **D.** Analysis of the distance traveled by different radius droplets obtained by modulating substrate concentration (10 mm and 100 mM) after 100 seconds. The data here reported represents a dataset of at least 3 different droplets.

### Droplets migration favours the formation of a simple metabolic pathway

In an origin of life scenario, the emergence of metabolic pathways is crucial for the evolution of living systems. From this evolutionary point of view, new metabolic pathways can emerge if different communities merge and mix their contents in a precise directional way [54, 55]. This scenario would not be favoured if the individuals cannot migrate or merge. As proof of concept of the importance of migration for reconstituting metabolic pathways, we selected the enzymes pyruvate kinase (PK) and L-lactate dehydrogenase (LDH) (partitioning reported in Fig. S2) to set up a simple two-enzyme pathway inside the droplets. The reaction product of PK is pyruvate (and ATP), which is the substrate of LDH that catalyze, in presence of NADH, the conversion of pyruvate to lactate and NAD^+^ (Fig. S3). Confocal microscopy images of droplets containing active unlabeled urease (along with either Alexa Fluor 488 labeled-PK or Alexa Fluor 594 labeled-LDH) show that multienzyme-containing droplets retain their migration feature when triggered by a chemical pH gradient (Fig. 5A and Supplementary Movie 10). This leads to the formation of a small and efficient linear pathway by merging two different droplets and mixing efficiently their contents (Fig. 5A). On the contrary, in absence of urease the droplets do not show migration and even when they stochastically merge, they display slower enzymes mixing compared to the active droplets. The slower mixing is due to the absence of Marangoni-induced flow and, thus, based on simple diffusion (Fig. 5B and Supplementary Movie 11). To understand how the reaction is influenced by the migration and merging of the droplets, we quantify the production of lactate (the final product of the two-step reaction). In the negative control (Fig. 5C, black dots and line), using separated droplets containing LDH or PK, the overall lactate production is slower compared to droplets containing LDH, PK and urease (Fig. 5C, purple dots and line). This result is in line with the fact that the pyruvate produced by the droplets with PK needs to diffuse out from the droplets and to enter into the droplets with partitioned LDH to produce lactate. To validate our hypothesis and the results just shown, we used droplets containing both urease and LDH, and droplets containing both urease and PK. Upon activation of urease and PK (with their substrates), the amount of lactate produced (Fig. 5C, yellow dots and line) is the same as the negative control in the first minutes, confirming the separate localization of the two enzymes in separate droplets. Interestingly, after 5-10 minutes, the production of lactate accelerates reaching the rate of the positive control, indicating that lactate production is increased by the co-localization of the two enzymes achieved through droplets migration and merging. This result is in agreement with the experiments obtained using the confocal microscope, which highlight droplets migration, as reported in Fig. 5A and B.

**Fig. 5.**
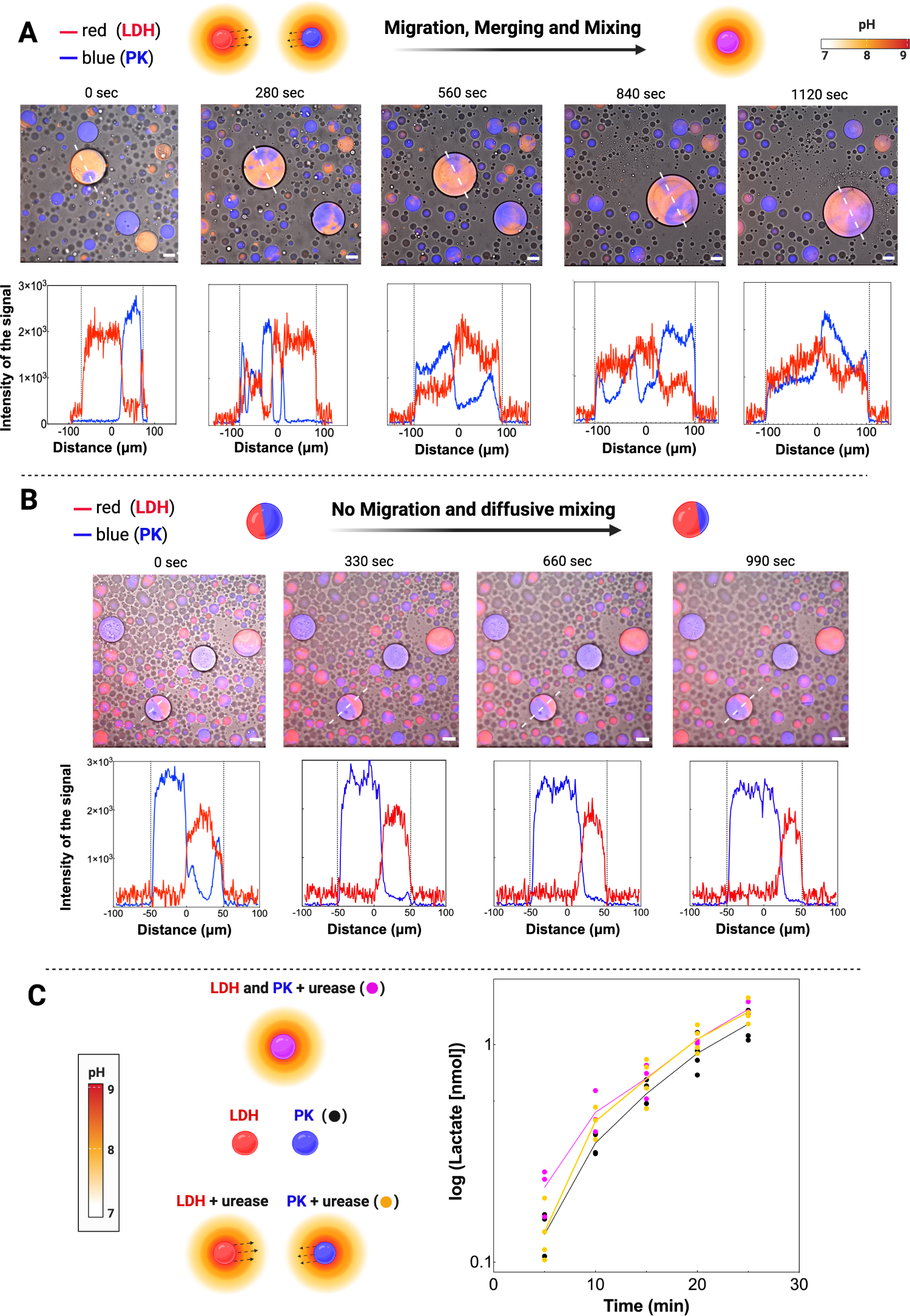
The formation of a short metabolic pathway is driven by the migration of the active droplets. **A.** Top to bottom: snapshots at different times of active droplets containing 1 *µ*M urease (not labeled) and 3.3 *µ*M Alexa Fluor 488 labeled-LDH (red channel) in presence of active droplets containing 1 *µ*M urease and 3.3 *µ*M Alexa Fluor 594 labeled-PK (blue channel) which display migration, merging and forced mixing mediated by the internal flow (when activated by adding 115 mM urea). The distribution of the different enzymes is highlighted by adjusting the colour channels separately. At the bottom are reported the intensity of the signal in blue and in red which are changing over time due to the mixing of the droplet content. Scale bar 50 *µ*m. **B.** Snapshots of active droplets at different times containing 3.3 *µ*M Alexa Fluor 488 labeled-LDH and 3.3 *µ*M Alexa Fluor 594 labeled-PK without urease. The distribution of the different enzymes is highlighted by adjusting the colour channels separately. At the bottom are reported the intensity of the signal in blue and in red which are changing over time due to the mixing of the droplet content. Scale bar 50 *µ*m. **C.** Evaluation of lactate produced by LDH using three different setups. Purple dots and line represent the positive control, based on active droplets (in presence of urea 115 mM, phosphoenolpyruvate 1 mM, NADH 2 mM and ADP 2 mM) containing 1 *µ*M urease, 3.3 *µ*M PK and 3.3 *µ*M LDH; Black dots and line represent the negative control of the experiment, based on active droplets (in presence of urea 115 mM phosphoenolpyruvate 1 mM, NADH 2 mM and ADP 2 mM) containing either 3.3 *µ*M LDH or 3.3 *µ*M PK, in separated droplets. In the last set up, yellow dots and line represent active droplets (in presence of urea 115 mM, phosphoenolpyruvate 1 mM, NADH 2 mM and ADP 2 mM) containing 1 *µ*M urease with 3.3 *µ*M LDH and droplets containing 1 *µ*M urease with 3.3 *µ*M PK. The solid line is the average value of each setup (*n ≥* 3), individual experimental points are shown (markers).

## Discussion

In the last decade, a wide pool of *in vitro* synthetic droplets systems have been generated. Spanning from membrane-embedded vesicles to membraneless droplets systems, they imposed themselves as key tool to study biochemical features of the cell and provide the basis for the creation of new materials. Reproducing cellular behavior *in vitro* is essential to understand how living cellular systems work and interact. However, mimicking cellular complexity *in vitro* is still challenging due to limitations of the current protocell systems.

In this study, we discovered that membraneless enzymatically-active droplets respond to a pH gradient in a chemotactic manner and that we can tune their migration by varying substrate and enzyme concentrations. These synthetic droplets mimic cytosolic protein crowding and resemble rudimental cellular features such as sensing and migration towards neighboring droplets by following the chemotactic gradient. Each droplet enzymatically produces ammonia, generating a pH gradient that forms a pH halo extending outside the droplets. The pH halo intensity and amplitude increase accordingly to the droplets radius (Fig. 2). Assuming that the pH halo is a fundamental prerequisite to “sense” the surrounding droplets, this analysis indicates that droplets (with radius *>* 30-35*µ*m) can chemically detect other droplets with long-range interactions mediated by the pH halo, with a sensing range that increase according to its amplitude. The pH change modifies the surface tension of the droplet [53], triggering a Marangoni-induced flow, driving the migrations of the droplets towards the neighboring ones. The migration velocity depends on the surface tension gradient along the droplet interface, with larger gradients resulting in higher migration speeds. Surface tension changes with the local pH value within a narrow range, approximately between pH 7.5 and 8.5. Small droplets (radius *<* 25-30 *µ*m) with low internal pH (pH *≤* 7.5) do not display migration, as the surface tension at their interface remains constant, and no significant Marangoni-induced flow occurs. In contrast, large droplets (radius *>* 35-40 *µ*m) show higher internal pH values, enabling sensing, migration and merging with neighboring droplets within 1-2 minutes of enzyme’s substrate addition (115 mM urea, Fig. 3A). Experimental data on droplet migration were confirmed through numerical simulations, showing that droplets move against the surface tension gradient along their interface, consistent with experimental observations. The flow directs the droplets towards low-surface tension regions (highest pH) generated by the neighboring active droplets. Internal pH is a key factor in determining droplet migration, influencing surface tension at the interface and extending the pH halo. Small droplets with low internal pH do not move, while larger droplets exhibit self-propelled migration, driven by their higher internal pH (Fig. 3B-E). We also observed small droplets moving within the halo produced by larger droplets (Fig. 3C). This behavior is explained by the imbalance in pH and surface tension along the interface of the smaller droplet, triggering Marangoni-induced migration towards the larger droplet. Our physics-aware explanation successfully reproduces these findings in numerical simulations, offering insight into the chemotactic migration and interaction of droplets. Moreover, we show how droplets migration speed can be tuned by modulating the enzyme activity inside the droplets. Indeed, by increasing either enzyme or substrate concentration, the droplets migration speed is enhanced accordingly (Fig. 4). Taken together, these data report a synthetic droplets system suitable for designing migrating systems with controllable migrating/delivery speed, based on the amount of enzyme/substrate provided.

The mechanisms we described here could represent a rudimental and ancient chemotactical sensing mean, which arose before the development of complex cellular machinery of chemotaxis. This phenomena could have played a key role in an origin of life scenario and could have driven the emergence of metabolic pathways. To support this hypothesis, we showed how the migration of droplets can facilitate the formation of a short metabolic pathway (Fig. 5). In this experiment, the enzyme triggered migration leads to (i) localization of the metabolic pathway in a single droplet, (ii) mixing due to the inner Marangoni-induced flow, and (iii) increased yield of the final product. Specifically, droplet migration speeds up the final product yield to the same levels of pre-localized control within the first 15 minutes of the reaction.

This study represents a significant stride in increasing the degree of complexity of synthetic droplets to emulate cellular features and effectvely integrates recent advances in this field [18, 19, 30, 35, 42, 43]. Notably, while previous research has reported random droplet motion driven by Marangoni flow [27, 56–59], it generally relied on the use of surfactants to initiate the Marangoni flow. In contrast, our study achieves droplet migration through enzymatic activity, which serves as the trigger for Marangoni flow. Additionally, it is important to highlight that our study does not merely demonstrate enzyme-generated random motion, as previously shown [60]. Instead, our work showcases droplet migration exclusively toward other droplets, resembling sensing and selective migration phenomena akin to cellular systems. This work represents an essential step for assessing biologically-relevant questions related to long-range interactions and chemotactical sensing. Furthermore, these results show the relevance of fluids mechanics in ubiquitous biological processes such as chemotactical sensing and migration, opening up interesting new perspectives as synthetic biological tool, as a drug delivery system and bio-inspired smart materials.

## Material and Methods

### Materials

Polyethylene glycol (PEG) 4000 Da (A16151) was purchased from Alpha-Aesar. Bovine serum albumin (BSA) (A7638), Potassium phospate dibasic trihydrate (60349), 3-(Trimethoxysilyl)propylmethacrylate (440159), 2-Hydroxy-4’-(2-hydroxyyethoxy)-2methylpropiophenone (410896), Poly(ethylene glycol) diacrylate (PEGDA) 700 Da (455008), PEGDA 4 kDa (907227), PEGDA 6 kDa (701963), PEGDA 10 kDa (729094), L-lactate dehydrogenase (LDH) from muscle rabbit (L1254), Pyruvate Kinase (PK) from muscle rabbit (P9136), Jack bean Urease (U4002), Peroxidase from horseradish (P8375), *β*-Nicotinamide adenine dinucleotide reduced disodium salt hydrate (NADH) (N8129), Phosphoenolpyruvate (860077), Lactate assay Kit (MAK064), Pyrydoxal 5’-phosphate (PLP) (P9255), glucose (G8270) and *β*-chloro-Lalanine (C9033) were purchased from Sigma-Aldrich. Potassium phosphate monobasic (42420) and trichloroacetic acid (TCA) (34603) was purchased from Nacalai Tesque. Alexa Fluor 594 and 488 Microscale Protein Labeling Kit (A30008 and A30006), Succinimidyl Ester SNARF-1 (S2280) were purchased from Thermo Fischer Scientific. Fluorescent polystyrene nanoparticles tracers of 0.2 *µ*m of diameter (FCDG003) was purchased from Bangs Laboratories.

### Glass coating slides preparation

Glass slides (VWR) were cleaned using water and ethanol and let to dry out for a few minutes. The clean glass slides were pre-treated with a 0.3% v/v 3-(trimethoxysilyl) propylmethacrylate dispersed in a 5% v/v ethanol-water mixture and then quenched adding pure ethanol. At this point a solution of the UV activated initiator 2-Hydroxy4’-(2-hydroxyyethoxy)-2methylpropiophenone at 1% w/w was prepared in water, and then added it to PEGDA 700 or 4000, 6000 or 10000 Da in a 80/20 initiator solution to PEGDA v/v ratio. 30 *µ*L of the PEGDA-initiator mixture were pipetted on a pretreated glass slide, and another clean, untreated glass slide (coated with an anti-rain film) was deposited on top. A thin layer of the PEGDA initiator mixture was allowed to spread via capillary action between the two glass slides to create a uniform coating. The glass slides were put under a UV lamp for 10 minutes and then stored underwater.

### Protein labeling

Bovine serum albumine, urease, cystalysin, pyruvate kinase and Lactate dehydrogenase were tagged as reported previously [53]. The reaction between SNARF-1 and BSA was carried out in water for 60 minutes at room temperature. All the tagged proteins were further purified using membrane dialysis to reduce the free probe in solution. The protein concentration of BSA, urease and Lactate dehydrogenase was measured as reported previously [53]. To determine the concentration pyruvate kinase we used *ε*_280_*_nm_* = 30410 M*^−^*^1^ cm*^−^*^1^ (https://web.expasy.org/protparam/) while the method for the quantification of cystalysin is reported in the Sec. Expression and purification of cystalysin. The degree of labeling of the proteins with the fluorescent probes was evaluated by following the manufacturer’s protocol (Microscale Protein Labeling Kit, ThermoFisher Scientific).

### Confocal imaging and movies analysis

All fluorescence and brightfield images were acquired and analyzed using a Spinning Disk Confocal (Nikon, Andor CSU) an LSM 880 Airyscan (Carl Zeiss Microscopy) microscopes with 63*×* oil immersion lens, 40*×*/1.30 PlanApo oil lens and 20*×* lens.

Confocal images were acquired for more than 3-5 independent experiments with similar results. The images collected were analyzed using ImageJ. All the movies reported have been recorded by using a Spinning Disk Confocal (Nikon, Andor CSU) using 20*×* lens. The Movie fps (frame per second) have been modified to convert the video timescale to 1 minute per second. The scale bar is equal to 100 *µ*m.

### pH Imaging and halo measurements

We quantified the pH variations inside the droplets using fluorescence ratiometric imaging as reported here [53]. After labelling BSA with SNARF-1 (with minor modifications from the previous protocol, by adding 100 *µ*M SNARF-1 as final concentration to the BSA solution), SNARF-1 labeled BSA was used to make droplets following the procedure reported in the section “Droplets preparation” in SI, with a final concentration of 1 *µ*M for urease, 33 *µ*M Cystalysin and 33 *µ*M Glucose oxidase inside the droplets. Note that the enzymes were not labeled with any fluorescent probe but the partitioning of the enzymes inside the droplets have been evaluated by a qualitative analysis of the intensity fluorescence in and out the droplets Fig. S2. However, the partitioning of urease by meauring its activity has been previously evaluated here [53]. To measure the halo around the active droplets free SNARF-1 probe was added to the supernatant phase at a final concentration of 50-100 *µ*M and added to the droplets along with the substrate. Briefly, the droplet solution was centrifuged for 30 seconds at 16900 g at room temperature and roughly more than half of the total supernatant was removed from the microcentrifuge tube (in this way, after resuspension the droplets are bigger). The droplets phase was then gently resuspended and 2 *µ*L was diluted in 18 *µ*L of free supernatant and added to the sample chamber. Then 180 *µ*L of supernatant containing the substrate and free SNARF-1 were added to the sample chamber and the measurement started. A confocal microscope LSM 880 Airyscan (Carl Zeiss Microscopy) has been used to image the pH inside and out-side the droplets. A 40*×*/1.30 PlanApo oil lens has been used for all the experiments.

Fluorescence imaging was performed exciting the dye at 561 nm and monitoring the pH-dependent emission spectral shifts simultaneously in two separate channels. The emission wavelengths selected were *λ_g_* = 585 nm and *λ_r_* = 640 nm. Only the emission fluorescence at *λ_r_* = 640 nm has been shown in (Fig. 2) to show the droplet halo.

### Pyruvate kinase and lactate dehydrogenase activity assay

Droplet suspensions with partitioned pyruvate kinase and l-lactate dehydrogenase were prepared as reported in the previous section, by adjusting the concentrations of inorganic ions and enzyme cofactors as follows: KCl 80 mM, MgCl_2_ 5 mM, ADP 2 mM and NADH 2 mM. The final concentration of enzymes inside the droplets were 3.3 *µ*M LDH, 3.3 *µ*M PK and 1 *µ*M urease. The experiments have been performed as follows. 15 *µ*L of droplets and supernatant phase was diluted in 735 *µ*L of supernatant phase (1:50 dilution) containing 1 mM phosphoenolpyruvate, 2 mM ADP, 2 mM NADH and 115 mM urea. At each timepoint, 45 *µ*L of reaction mix was quenched using 5 *µ*L of 100% Trichloroacetic acid (TCA) (dilution 1:10 v/v) and stored in ice. Then, the quenched samples were centrifuged for 10 minutes at 16000 g at RT (25*^◦^*C). The amount of lactate in the reaction mix was measured with the Lactate Assay Kit (colorimetric assay) by reading the absorbance using Multiskan SkyHigh plate reader (Thermo Fisher Scientific). Specifically, 5 *µ*L of quenched samples were added to 45 *µ*L of Lactate Assay Buffer (provided in Lactate Assay Kit) into a 96 well plate. At this point, 50 *µ*L of solution (Lactate Probe + Lactate Enzyme Mix + Lactate Assay Buffer) was added to the stopped reaction mix and incubated for 30 minutes protected from light. Absorbance was read at 570 nm. Lactate from the Lactate Assay Kit was used to determine the nmoles of lactate present in the wells through the calibration curve. The data were analyzed using GraphPad Prism 10 and at least three independent experiments were performed.

### Numerical method

To simulate the dynamics of the droplets we solved the incompressible Navier-Stokes equations (conservation of momentum and incompressibility constraint) coupled to the volume of fluid transport equation and to the transport equation of the concentration field determining the local pH value. The droplets were identified using a color function, the volume of fluid variable, which distinguishes the droplets from the supernatant. The multi-dimensional tangent of hyperbola for interface capturing (MTHINC) method [61, 62] was used within the volume of fluid framework to improve the accuracy in the transport of the interface and to reduce parasitic currents at the interface. The surface tension of the interface of the droplets depends on the local pH field [53]; the change in surface tension as a function of the local pH is reported in Fig. S8. An advection-diffusion equation was used to describe the transport of the concentration field determining the local value of the pH. A size-dependent source term is active within the droplets to model the chemical reaction occurring inside the droplets and increasing the pH value.

The system of equation was discretized in space using a second-order central finite difference scheme on a Cartesian, uniform grid. Time-advancement is performed using the second-order explicit Adams-Bashforth scheme. We used the in-house flow solver Fujin (https://groups.oist.jp/cffu/code), used and validated for a variety of problems [62–65]. Further details on the numerical method can be found in the Supplementary information.

### Statistics and Reproducibility

All the experimental data presented in this work have been performed in triplicate at least. The authors were not blinded during outcome assessment or in the experimental design.

## Data availability

The dataset generated during the study are available from the corresponding authors on reasonable request.

## Code availability

The code used for the present research is a standard direct numerical simulation solver for the Navier–Stokes equations. Full details of the code used for the numerical simulations are provided in the Methods section, in the Supplementary information, and in the references therein.

## Author Contributions

M.D., A.M. and P.L. conceived the project. M.D., A.B. and P.L designed the experiments. M.D. and A.B. performed all the experiments. M.D. A.B. and P.L. analyzed the experimental data. G.S., A.M. and M.E.R. wrote the code, performed the simulations and analyzed the numerical data. P.L., M.D., A.B. G.S. and M.E.R. wrote the manuscript. This project was supervised by P.L.

## Acknowledgements

The research was supported by the Okinawa Institute of Science and Technology Graduate University (OIST) with subsidy funding to P.L. and M.E.R. from the Cabinet Office, Government of Japan. M.D. thanks the financial support from Japan Society for the Promotion of Science (JSPS) for the Kakenhi Early Career Scientist N. 22K15065. P.L. thanks the financial support from Takeda Grant. We are grateful for the help provided by imaging section at OIST; in particular we thank Paolo Barzaghi for the support with the confocal microscopes. We also thank Stefano Pascarelli for the suggestions on the halo analysis, Mohamed Abdelgawad for the help with the droplet tracking analysis, Prof. Amy Shen for the initial discussion on the coating polymers and Prof. Barbara Cellini for providing the Cystalysin expression vector. The authors would also like to thank Ms Amy Gooch for critical reading. M.E.R. and G.S. acknowledge the computational resources provided by the Scientific Computing Section of the Research Support Division at OIST and the computational resources provided by HPCI under the grant hp220100. The images of the present work were prepared using Biorender.com.

