## Supplementary Information for "Chemotactic interactions drive migration of membraneless active droplets"

### Supplementary information: Chemotactic interactions drive migration of membraneless active droplets

#### Expression and purification of cystalysin

We adopt the conditions used for expression of the wild-type cystalysin reported in previous works [1], with a minor modification. Briefly, the expression of the protein has been performed using BL21 expression cells instead of DH5$\alpha$. The protein concentration in all cystalysin samples was determined by absorbance spectroscopy [2] using $\varepsilon_{280nm} = 127000$ M$^{-1}$ cm$^{-1}$. The PLP content of the enzyme was determined by releasing the coenzyme in 0.1 M NaOH and by using $\varepsilon_{388nm} = 6,600$ M$^{-1}$ cm$^{-1}$ [3].

#### Droplet preparation

A polyethylene glycol (PEG) stock solution at 600-650 mg/mL ( 60% w/w) was prepared mixing PEG 4000 Da and Milli-Q water. A Bovine Serum Albumin (BSA) stock solution with a target concentration of 5-7 mM was prepared in Milli-Q water. The concentration of BSA was confirmed by measuring the absorption intensity at the 280

---

[†]These authors contributed equally to this work

nm peak with a UV-vis spectrophotometer (UV-1900 UV-Vis Spectrophotometer, Shimadzu), assuming $\varepsilon_{280nm} = 43,824$ M$^{-1}$ cm$^{-1}$ (https://web.expasy.org/protparam/). The droplet suspensions were prepared mixing appropriate amounts of PEG and BSA stock along with other components. In detail, the reagents have been added and mixed following a specific order: first we added the appropriate volume of buffer KP 0.2 M pH 7.0 and small molecules (enzymes, cofactors, salts, fluorescent nanoparticles and substrates at the appropriate concentration); after mixing, we added BSA at the final concentration of 447 $\mu$M. After mixing, as last component of the system, PEG was slowly added (23 % w/w final) and the solution has been carefully mixed, and the composition of the droplets was checked qualitatively at the confocal microscope. If required, the supernatant phase was isolated centrifuging the droplet suspension for 30 minutes at 16,900 g at 25°C and extracting the supernatant phase with a pipette (without withdrawing the droplet phase).

### Evaluation of enzymes partitioning through enzymatic activity measurement

The partitioning of the enzymes selected for this study (L-lactate dehydrogenase (LDH), Urease, Piruvate kinase (PK), Glucose oxidase and Cystalysin) was evaluated by measuring the activity of the enzyme in both the supernatant and the droplets phase, after they were separated by centrifugation. Specifically, 1 mL of droplets suspension containing a fixed enzyme concentration was prepared and centrifuged for 30 minutes at 16900 g at room temperature. At this point, the supernatant phase was separated from the droplets phase into another tube. Both phases were diluted 1:4 (using potassium phosphate buffer 0.1 M pH 7.0.) and the enzymatic activity of the single phases was evaluated. Enzymatic activity of Urease and LDH was tested as reported here [4]. Enzymatic activity of Glucose Oxidase, Pyruvate Kinase and Cystalysin was measured as follows:

- The production of hydrogen peroxide catalyzed by glucose oxidase (derived from the oxidation of a saturating concentration of D-glucose over time) was detected by monitoring the signal at 460 nm given by the oxidation of the cromogenic substrate o-dianisidine catalyzed by the Peroxidase. The droplets were generated containing 2-3 $\mu$M Glucose Oxidase and 20-30 $\mu$M Peroxidase (10 times higher concentration in order to completely convert the hydrogen peroxide produced by Glucose Oxidase). The signal was measured in continuous during the reaction by reading the absorbance using Multiskan SkyHigh plate reader (Thermo Fisher Scientific). The buffer used for the activity measurements was potassium phosphate 0.1 M pH 7.0.
- The production of pyruvate catalyzed from pyruvate kinase and cystalysin was measured with the same method. Briefly, 3-5 $\mu$M enzyme (inside the droplets) and a saturating concentration of the substrates has been used to test their partitioning in the droplet phase (1 mM phosphoenolpyruvate and 2 mM beta-chloro-L-alanine). At each time point, the reaction mix was stopped using trichloroacetic acid (TCA) (10% final, 45 $\mu$L reaction mix + 5 $\mu$L of TCA 100%) and stored in ice. The stopped samples were then centrifuged for 10 minutes at 16900 g at room temperature and the amount of pyruvate produced was measured using a spectrophotometric assay

coupled with LDH following the NADH signal at 340 nm, using a Multiskan Go plate reader (Thermo Fisher Scientific). The buffer used for the activity measurements was potassium phosphate 0.1 M pH 7.0.

### Numerical method

The set of equations defining the dynamics of the system is made of the momentum conservation equation, Eq. S1, the incompressibility constraint, Eq. S2, the volume of fluid transport equation, Eq. S3, and the transport equation for the scalar field $c$ determining the local pH value, Eq. S4.

$$\rho \frac{\partial \mathbf{u}}{\partial t} + \rho \mathbf{u} \cdot \nabla \mathbf{u} = -\nabla p + \nabla \cdot \left[\mu \left(\nabla \mathbf{u} + \mathbf{u}^T\right)\right] + \nabla \cdot (\tau_c f_\sigma), \tag{S1}$$

$$\nabla \cdot \mathbf{u} = 0, \tag{S2}$$

$$\frac{\partial \phi}{\partial t} + \mathbf{u} \cdot \nabla \phi = 0, \tag{S3}$$

$$\frac{\partial c}{\partial t} + \mathbf{u} \cdot \nabla c = \nabla \cdot (D \nabla c) + \phi \dot{Q} (\bar{c} - c). \tag{S4}$$

In the above equations, $\rho$ and $\mu$ are the density and dynamic viscosity of the fluid. In the numerical simulations we chose to use the same values of density and viscosity for both the supernatant and the droplets, as we observed that different material properties among the two phases do not significantly alter the general dynamics of the Marangoni-induced motion. $\mathbf{u}$ is the fluid velocity, $p$ the pressure, $\tau_c$ the Korteweg tensor, $f_\sigma$ the surface tension equation of state, $\phi$ the volume of fluid variable, $c$ the local concentration of a scalar field determining the local value of pH, and $D$ its diffusion coefficient. A source term is active inside the droplets (i.e. $\phi = 1$) with magnitude imposed by $\dot{Q}$; the source term forces the concentration inside the droplets to a set value, $\bar{c}$, reported in table 1 (labeled as internal pH$^*$). Due to the explicit time advancement, the value of the diffusion coefficient is lower than the experimental value, as indicated by the smaller width of the halo around the droplets in the numerical simulations. For this reason, in our numerical simulations we opted for a linear relation between the local concentration field $c$ and the dimensionless pH value, $c = \text{pH}^* = (\text{pH} - \text{pH}_{min})/(\text{pH}_{max} - \text{pH}_{min})$, such that the resulting pH halo extends further from the interface of the droplet.

We do not simulate the reaction of urea and urease, and instead simulate directly the concentration field that causes the pH distribution. Experimental observations show that the timescale of the urea-urease reaction is much shorter than the timescale of the droplet motion, hence the reaction can be well approximated as nearly instantaneous. The production of ions from the reaction, causing an increase in the pH, is here modelled using a source term. The simulation of the urea-urease reaction would require the use of additional models, variables and free parameters that must be chosen, tuned, and further validated. Simulating the urea-urease reaction would also significantly increase the computational cost of the simulations (additional variables, coupled equations and stricter constraints on the time-advancement due to the fast dynamics), while adding little to the overall physical insight.

The Korteweg tensor $\tau_c = (\mathbf{n} \cdot \mathbf{n})\mathbf{I} - \mathbf{n} \otimes \mathbf{n}$ [5] characterizes the interface ($\mathbf{n} = \nabla\phi/|\nabla\phi|$ is the unit-length vector normal to the interface) and is used to compute the surface tension forces generated by the interface. When surface tension is uniform along the interface (i.e. $f_\sigma$ uniform along the interface), the surface tension forces have only a component normal to the interface, the capillary stresses $f_\sigma \nabla \cdot \tau_c$. Conversely, when surface tension changes along the interface, an additional tangential contribution appears, the Marangoni stresses $\tau_c \cdot \nabla f_\sigma$. This latter component is crucial in driving the motility of the droplets: Marangoni stresses, tangential to the interface, generate a flow inside and outside the droplets, which causes the droplets to self propel. Marangoni stresses are proportional to the gradient of surface tension; experimental measurements [4] showed a reduction of surface tension for increasing values of pH in the range of pH between 7.2 and 8.4, hence Marangoni stresses are directed against the pH gradient (from high pH – low surface tension regions towards low pH – high surface tension regions). A single, isolated droplet generates a pH distribution which is uniform along its interface and thus does not generate any tangential stresses. On the other hand, the interaction of a droplet with the halo formed by another droplet generates a non-uniform surface tension distribution along the interface, and thus Marangoni stresses. Marangoni stresses drive the droplets towards each other, as the local value of pH is higher (i.e. lower surface tension) on the side where the droplets face each other.

#### Surface tension equation of state

The surface tension equation of state that has been selected for the numerical simulations is reported in Fig. S8. This equation of state has been selected to approximate the experimental measurements [4], which are also reported in the inset in the same figure. According to the experimental measurements [4], the surface tension is roughly constant at low and high values of pH and steeply decreases at intermediate pH values. The change in surface tension is indeed limited in the range of pH between pH$\approx 7.5$ and pH$\approx 8.2$. The proposed model reproduces this behavior, with a roughly constant (high) surface tension at low values of pH, a steep decrease in surface tension at intermediate pH values, followed by a roughly constant (low) surface tension at high pH values. We model the surface tension equation of state using a hyperbolic tangent function: $\sigma/\sigma_0 = \alpha \tanh\left[\gamma(\text{pH}^* - \text{pH}^*_0)\right] + \beta$, where $\text{pH}^*$ is the non-dimensional pH value $\text{pH}^* = (\text{pH} - \text{pH}_{min})/(\text{pH}_{max} - \text{pH}_{min})$. The surface tension at $\text{pH}^* = 0$ is $\sigma_0$ and the maximum surface tension gradient occurs at $\text{pH}^*_0 = 0.7$. The parameter $\gamma = 10$ sets the width of the region characterized by a steep decrease of the surface tension. The numerical constants $\alpha = -0.451116$ and $\beta = 0.548885$ set the range and the minimum value of surface tension; here the surface tension at the maximum surface tension concentration $\text{pH}^* = 1$ is equal to 20% of the reference value $\sigma_0$.

### Computational setup

Three-dimensional numerical simulations are performed in a regime similar to that of the experiments. The length of the domain $L_x$ and the characteristic velocity of the droplets $u$ are used as length and velocity scales. The Reynolds number (ratio of inertial over viscous contributions) is very small, $Re_0 = \rho u L_x/\mu = 10^{-2}$, indicating the dominance of viscous contribution over inertial ones. The Péclet number (ratio of advective transport over diffusive transport) of the concentration field is $Pe_0 = uL_x/D = 10^{-2}$, in agreement with the fast diffusion (compared to advection) observed in the experiments. The capillary number (ratio of viscous contributions over surface tension forces) is $Ca = \mu u/\sigma_0 = 10^{-4}$, indicating the strong effect of surface tension forces. The source term $\dot{Q}$ is set to $\dot{Q} = 10^5 u/L_x$ in order to guarantee an almost constant concentration inside the droplets, in agreement with the experiments. With this set of parameter, we obtain a Marangoni number of the order $\mathcal{O}(10)$. The Marangoni number, $Ma = \Delta\sigma L_x/\mu D$, is the ratio of Marangoni-induced transport over the diffusivity of the pH (i.e., of the concentration field).

The grid resolution is chosen to properly simulate the dynamics of the system and, in particular, interfacial phenomena, with the smallest droplet discretized with about 35 grid points per diameter. Numerical simulations performed on a twice-refined grid (in all directions, $\times 8$ number of grid points) showed negligible changes in the trajectory, velocity and dynamics of the Marangoni-driven droplets; further details on the grid independence tests are provided in the next section. The time step was selected to verify both the advective and diffusive Courant-Friedrichs-Lewy conditions: $\Delta t u/\Delta x = 8 \cdot 10^{-7} < 1$ and $\Delta t \mu/\Delta x^2 = 1.25 \cdot 10^{-3} < 1$. Given the low Reynolds number of the numerical simulations, the diffusive condition is the most critical. Surface tension forces further increases the constraints; hence we chose $\Delta t\mu/\Delta x^2 \ll 1$ to prevent the numerical solution from diverging. Table 1 summarizes all the numerical simulations performed and the computational setup adopted.

The computational domain is a closed box bounded by solid walls in all directions. No-flux boundary conditions are imposed on the velocity component normal to the domain boundaries and no-slip boundary conditions on the tangential component. No-flux boundary conditions are imposed to the volume of fluid as well (i.e. the droplets cannot leave the closed computational box), while we impose a far-field value at the domain boundary for the concentration field.

All cases start with the fluid at rest (zero velocity); the droplets are initialized with an internal pH value which depends on their size, as observed in the experiments. No halo is present at the beginning of the simulation: the chemical reaction has just started inside the droplets and the pH has yet to diffuse. Once the simulation starts, the development of the halo is extremely fast compared to the Marangoni-induced flow dynamics, as expected in the low-Marangoni number regime.

#### Grid independence tests

We performed grid independence tests on the case MDMD, see Tab. 1 for the list of simulations. Two cases were run: a coarser and a finer grid case. The grid resolution was increased in all directions by a factor two for the fine-grid case ($8\times$ the number of points) and reduced in all directions by a factor two for the coarse-grid case ($0.125\times$ the number of points). We show the horizontal (along the center-to-center axis) motion of the two droplets of the MDMD cases for all grid resolutions in Fig. S9. The figure shows the horizontal position of the center of each droplet over time; the droplets start to move towards each other till they merge at about $t = 0.08$. The grid resolution for the coarser case is sufficient to capture the dynamics of the droplets and already provides satisfactory results; refining the grid leads to a slightly earlier merging with overall similar dynamics. Negligible differences are observed among the fine-grid case and the standard grid we used in our analyses: the trajectory of the droplets fall on top of each other and the merging occurs at very similar times. The fine-grid case has a slightly later merging: this is easily explained as the thin liquid film in between the two droplets right before merging occurs is resolved with a higher grid resolution. This is typical of interface capturing methods, such as the volume of fluid, which use an Eulerian color function to identify the position of the droplets and of the interface.

### Supplementary figures

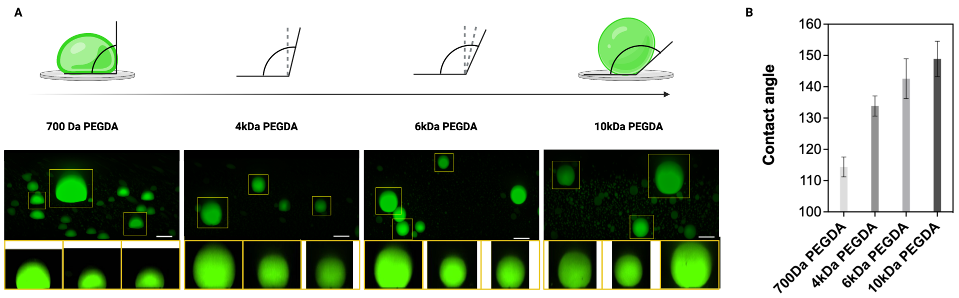


**Fig. S1 Contact angle analysis of sessile droplets on different PEGDA glass coating. A.** (from left to the right) z-stack images of Alexa Fluor 488-labeled BSA droplets using 700 Da, 4 kDa, 6 kDa and 10 kDa PEGDA glass coating. The droplets selected for the contact angle analysis have been highlighted. The side view images of the droplets used for the analysis are reported below each z-stack picture. Scale bar 20 $\mu$m. **B.** Contact angle $\theta$ analysis of the selected PEGDA glass coating. The results represents the average value of the analysis performed in triplicate on the droplets shown in panel A; error bars correspond to $\pm$SD. The analyzed droplets have a radius spanning from 8 to 30 $\mu$m.

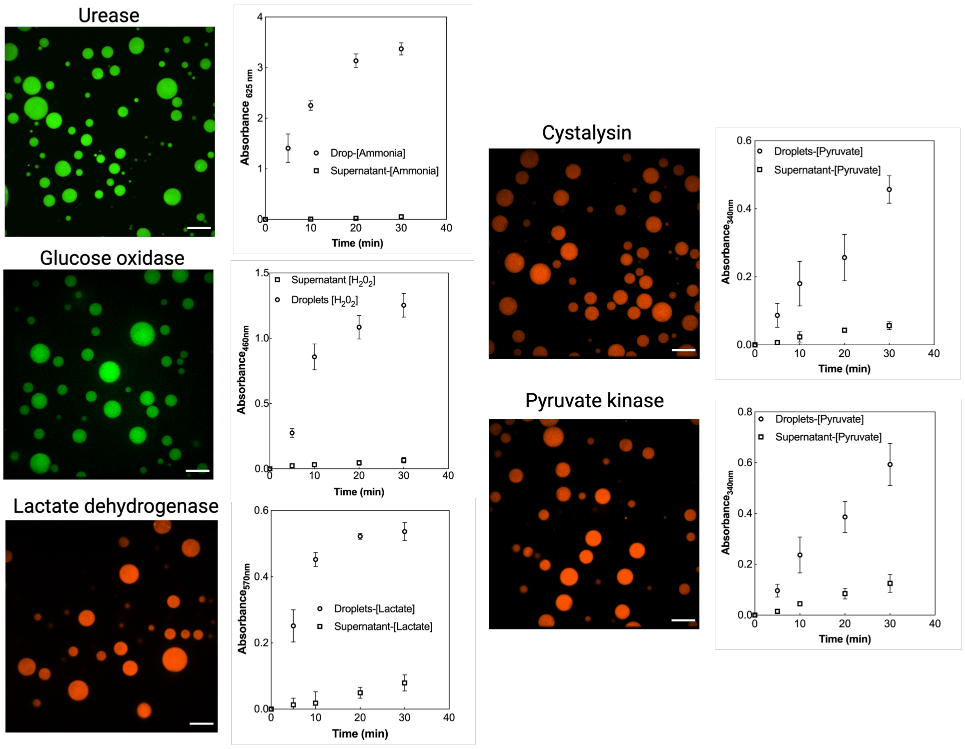


**Fig. S2 Evaluation of enzymes partitioning at the confocal microscope and by measuring the enzymatic activity.** Confocal microscope images of Alexa Fluor 594-labeled Cystalysin (33 $\mu$M), Alexa Fluor 488-labeled Glucose oxidase (33 $\mu$M), Alexa Fluor 594-labeled Pyruvate kinase (2 $\mu$M), Alexa Fluor 488-labeled Urease (1.5 $\mu$M) and Alexa Fluor 594-labeled Lactate dehydrogenase (33 $\mu$M). The droplets phase and the supernatant phase have been prepared as reported in the "Droplet preparation" section of the Supplementary Information. Scale bar is 25 $\mu$m. Enzymatic activity is reported in both droplet phase and in the supernatant phase only. The enzymes concentration within the droplets was 0.1-5 $\mu$M and the set-up of the enzymatic assay is reported in the "Evaluation of enzymes partitioning through enzymatic activity measurement" section of the Supplementary Information.

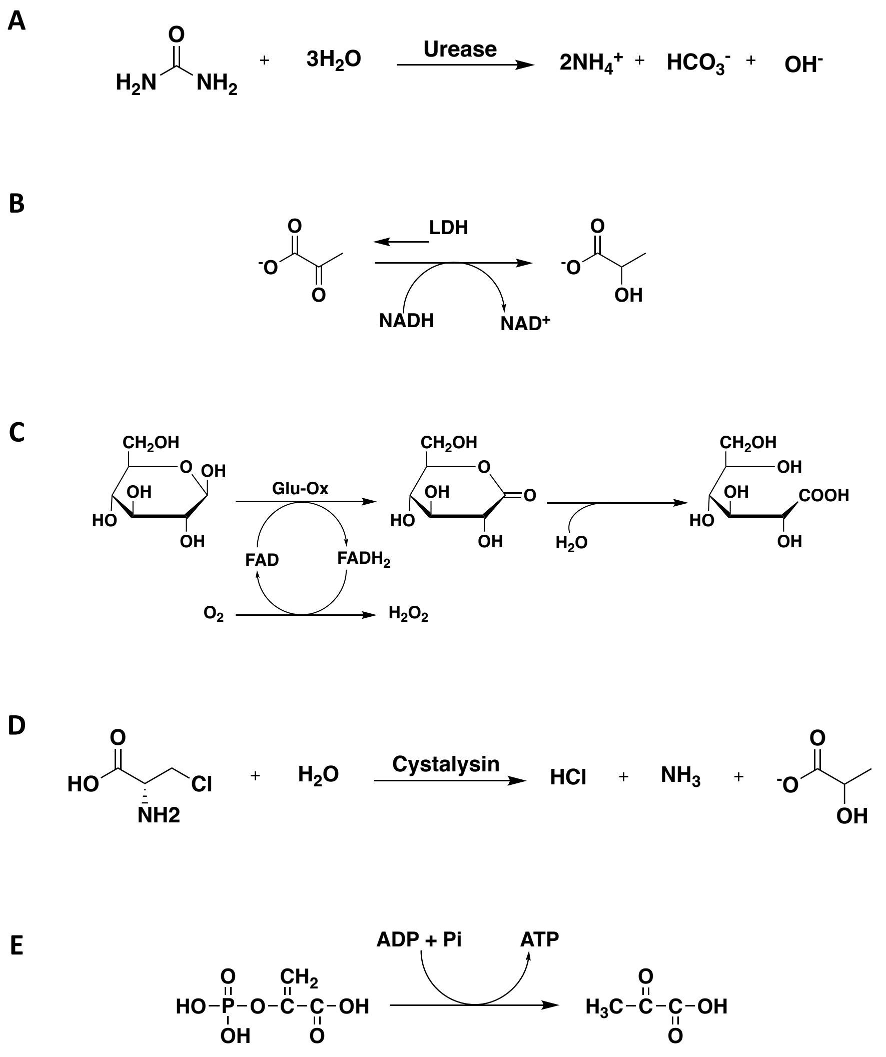


**Fig. S3 Reaction schemes catalyzed by the selected enzymes of this study.** Scheme of the chemical reactions performed by Urease **(A)**, Glucose Oxidase **(B)**, Lactate Dehydrogenase **(C)**, Cystalysin **(D)** and Pyruvate Kinase **(E)**.

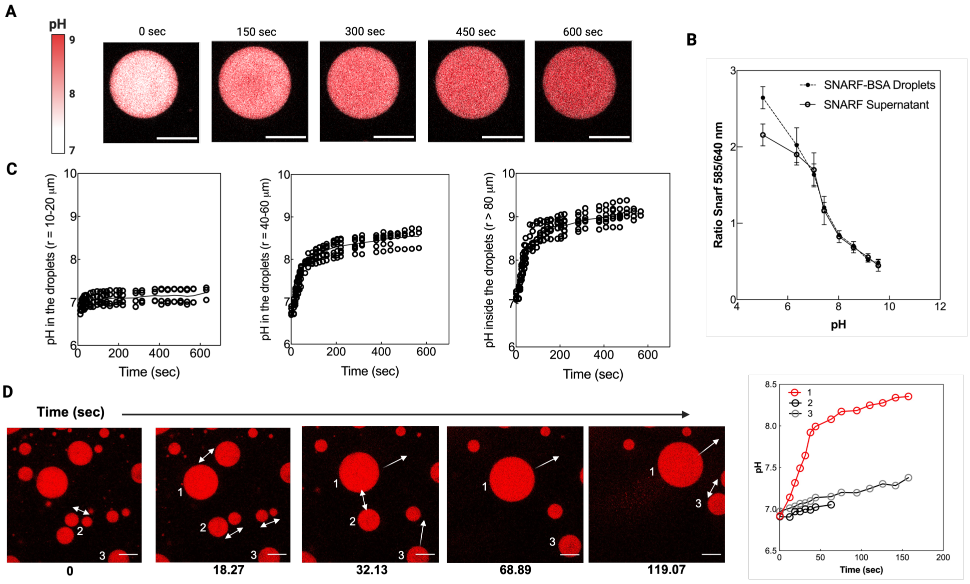


**Fig. S4 Internal pH change of active droplets loaded with urease and internal pH change of moving droplets of different size. (A)** pH change measured by using the pH sensitive probe SNARF-1 inside the droplets over time. **(B)** Calibration curve based on ratiometric analysis of SNARF-1 inside and outside the droplets. **(C)** Internal pH change graphs over time as a function of droplets size. **(D)** Image sequence of a big primary droplet (numbered as 1) and two secondary smaller droplets (numbered as 2 and 3) moving following a chemical gradient over time. The overall urease concentration inside the droplets was 1.5 $\mu$M and the substrate concentration was 115 mM. The BSA inside the droplets is labeled with SNARF-1 (red channel, emission wavelength 640 nm) to measure the pH change which values over time are reported for the three main droplets analyzed. Scale bars are 50 $\mu$m.

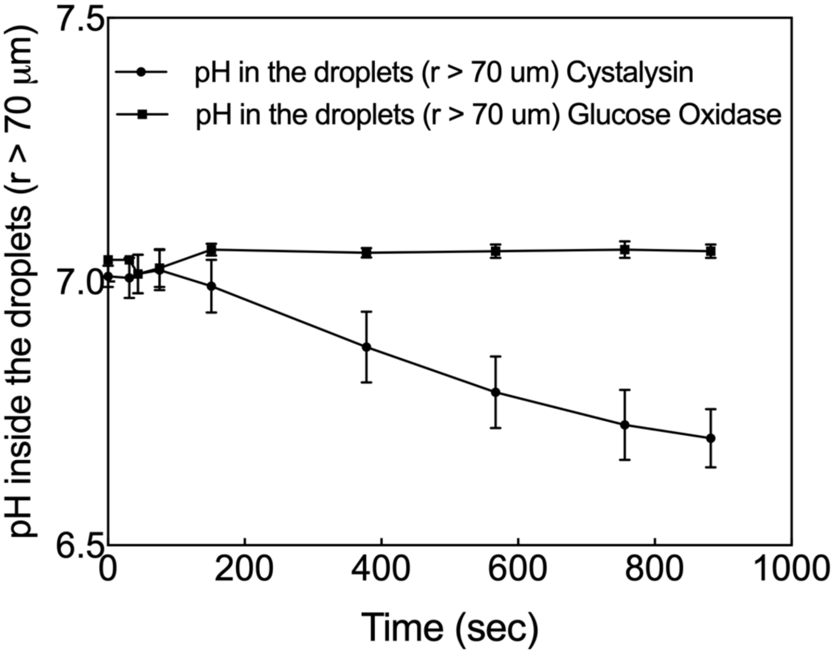


**Fig. S5 Evaluation of pH change over time in Cystalysin and Glucose Oxidase containing droplets.** The pH measurement was performed using droplets with radius $> 70$ $\mu$m containing cystalysin 33 $\mu$M and Glucose Oxidase 33 $\mu$M (inside the droplets) using BSA labeled with SNARF-1. The experiments using droplets containing-cystasylin have been performed in presence of 20 mM $\beta$-chloro-L-alanine and 100 $\mu$M PLP while those using droplets containing-glucose oxidase in presence of 50 mM glucose. Data are represented as mean values $\pm$SD from three independent droplets.

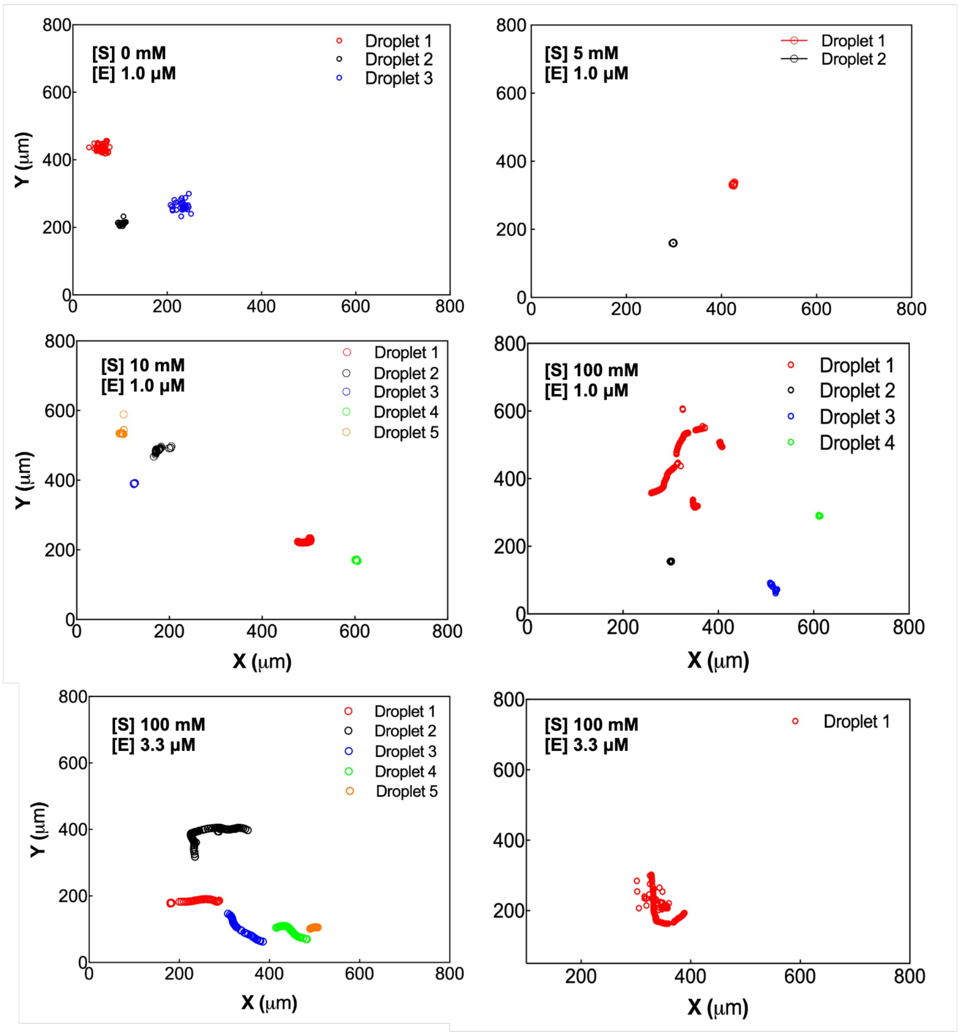


**Fig. S6 Trajectories (X,Y) calculated over time of droplets by modulating substrate and enzyme concentration.** Droplet trajectory analysis performed by tracking the center of each droplet obtained by varying substrate concentration from 0 to 200 mM urea while enzyme urease concentration was 1 $\mu$M or 3.3 $\mu$M. The droplets' trajectory has been analyzed using MATLAB software and every marker is drawn every $\Delta T = 2$ sec. The droplets sizes analyzed here are larger than 35 $\mu$m.

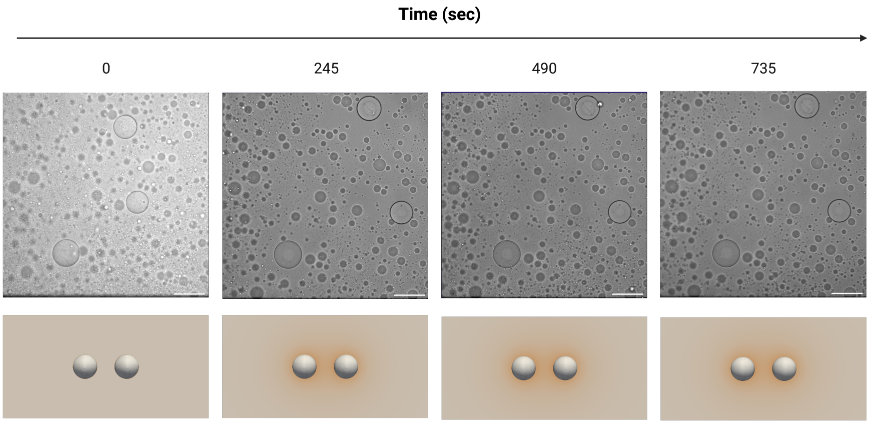


**Fig. S7 Small radius active droplets do not display migration.** Droplets with radius < 40 $\mu$m do not show migration. Time series of active droplets containing 1 $\mu$M urease in presence of 115 mM urea. The screenshots have been obtained from the supplementary movie 16. These data are supported by computational simulation of case SDSD (below). Scale bar 100 $\mu$m.

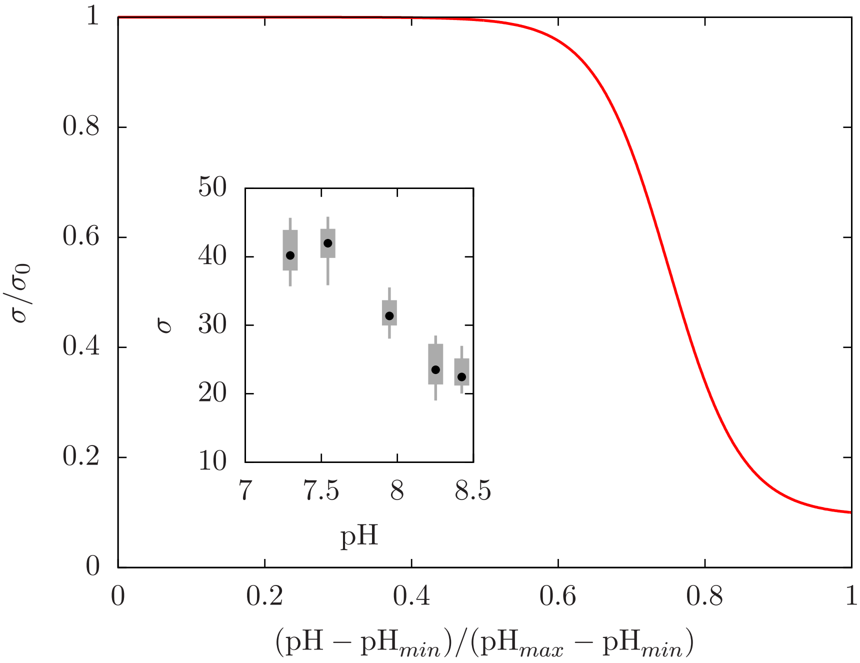


**Fig. S8 Surface tension equation of state as function of the dimensionless pH.** Experimental data [4] are reported in the inset.

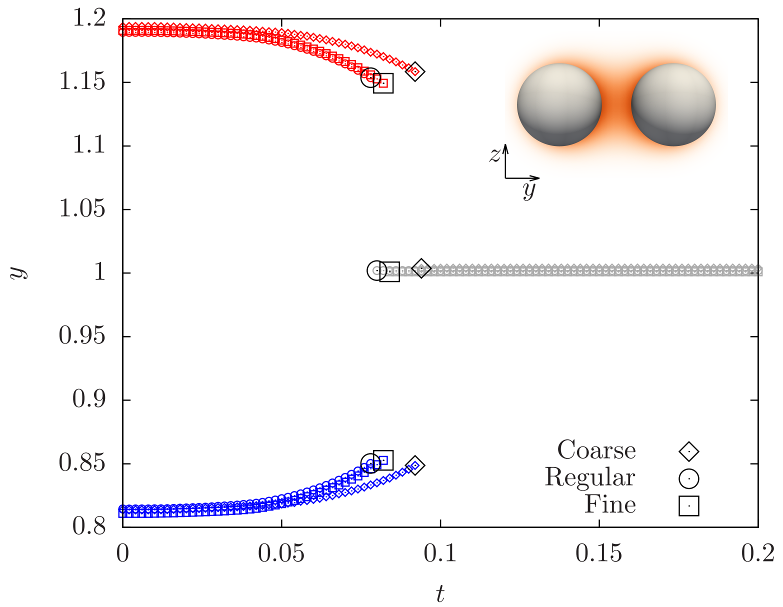


**Fig. S9 Grid independence test.** The horizontal position of each droplet over time is shown for the MDMD cases, all grid resolutions. The left-side droplet is reported in blue, the right-side droplet in red and the newly-formed droplet generated by the merging in grey; the merging of the droplet is identified by a larger black marker.

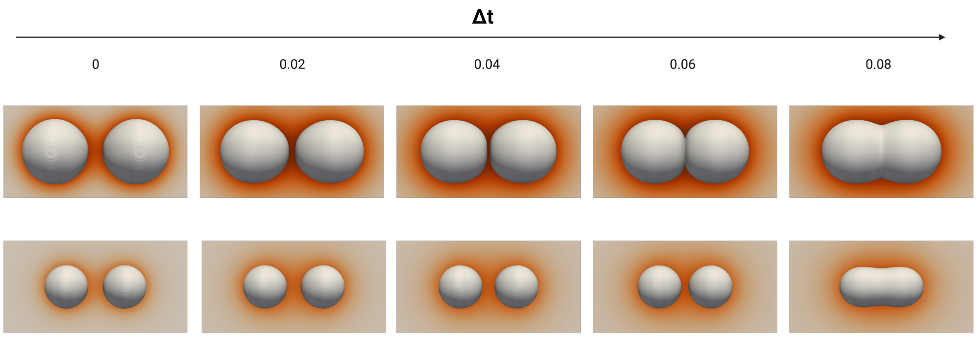


**Fig. S10 Increasing droplet radius speeds up its migration.** Time series of cases LDLD (top row) and MDMD (bottom row) from supplementary movie 15 and 12, respectively. Same scale is used for all time sequences.

**Table 1  List of simulations performed.** The size of the computational domain, the grid resolution, and the number, initial position and radius of the droplets is reported for each case. For the two-drop cases the acronym identifying the case indicates the size of the droplet: SD (small droplet, $R = 0.07L_x$), MD (medium droplet, $R = 0.14L_x$) and LD (large droplet, $R = 0.21L_x$).

| Case | Domain size | | | Computational grid | | | Number of droplets | Initial position | Radius | Internal pH* |
|---|---|---|---|---|---|---|---|---|---|---|
| | $L_x$ | $L_y$ | $L_z$ | $N_x$ | $N_y$ | $N_z$ | | | | |
| SDLD | 1.0 | 2.0 | 1.0 | 256 | 512 | 256 | 2 | $x_c = 0.5$ $y_c = 0.74$ $z_c = 0.5$ | 0.21 | 1.0 |
| | | | | | | | | $x_c = 0.5$ $y_c = 1.12$ $z_c = 0.5$ | 0.07 | 0.5 |
| MDMD | 1.0 | 2.0 | 1.0 | 256 | 512 | 256 | 2 | $x_c = 0.5$ $y_c = 0.81$ $z_c = 0.5$ | 0.14 | 0.75 |
| | | | | | | | | $x_c = 0.5$ $y_c = 1.19$ $z_c = 0.5$ | 0.14 | 0.75 |
| MDMD coarse grid | 1.0 | 2.0 | 1.0 | 128 | 256 | 128 | 2 | $x_c = 0.5$ $y_c = 0.81$ $z_c = 0.5$ | 0.14 | 0.75 |
| | | | | | | | | $x_c = 0.5$ $y_c = 1.19$ $z_c = 0.5$ | 0.14 | 0.75 |
| MDMD fine grid | 1.0 | 2.0 | 1.0 | 512 | 1024 | 512 | 2 | $x_c = 0.5$ $y_c = 0.81$ $z_c = 0.5$ | 0.14 | 0.75 |
| | | | | | | | | $x_c = 0.5$ $y_c = 1.19$ $z_c = 0.5$ | 0.14 | 0.75 |
| SDSD | 1.0 | 2.0 | 1.0 | 256 | 512 | 256 | 2 | $x_c = 0.5$ $y_c = 0.88$ $z_c = 0.5$ | 0.07 | 0.5 |
| | | | | | | | | $x_c = 0.5$ $y_c = 1.12$ $z_c = 0.5$ | 0.07 | 0.5 |
| 4 drops | 1.0 | 2.0 | 2.0 | 256 | 512 | 512 | 4 | $x_c = 0.5$ $y_c = 0.69$ $z_c = 0.74$ | 0.21 | 1.0 |
| | | | | | | | | $x_c = 0.5$ $y_c = 0.69$ $z_c = 1.12$ | 0.07 | 0.5 |
| | | | | | | | | $x_c = 0.5$ $y_c = 1.2$ $z_c = 0.9$ | 0.07 | 0.5 |
| | | | | | | | | $x_c = 0.5$ $y_c = 1.2$ $z_c = 0.1$ | 0.07 | 0.5 |
| LDLD | 1.0 | 2.0 | 1.0 | 256 | 512 | 256 | 2 | $x_c = 0.5$ $y_c = 0.74$ $z_c = 0.5$ | 0.21 | 1.0 |
| | | | | | | | | $x_c = 0.5$ $y_c = 1.26$ $z_c = 0.5$ | 0.21 | 1.0 |

### Supplementary movies

- **Supplementary Movie 1. Marangoni related flow and the droplet migration aim towards the same direction in a merging event between two active droplets of different sizes.** This movie report the Supplementary movie 8 with highlighted fluorescent nanoparticles (diluted 1:10000) to show Marangoni related flow inside the dropelts. The Movie fps have been modified to convert the video timescale to 1 minute per second. Scale bar 100 $\mu$m.
- **Supplementary Movie 2. Marangoni related flow and the droplet migration aim towards the same direction in multiple merging events between multiple active droplets of different sizes.** In the movie, 20 $\mu$L of diluted droplets phase (diluted 1:10, 2 $\mu$L of droplets phase + 18 $\mu$L of supernatant phase) containing 1 $\mu$M urease and fluorescen nanoparticles (diluted 1:10000), have been mixed with 180 $\mu$L of supernatant phase containing 115 mM Urea (final concentration). The Movie fps have been modified to convert the video timescale to 1 minute per second. Scale bar 100 $\mu$m.
- **Supplementary Movie 3. Droplets do not show migration in absence of urease's substrate.** In the movie, 20 $\mu$L of diluted droplets phase containing 1 $\mu$M urease (diluted 1:10, 2 $\mu$L droplets phase + 18 $\mu$L of supernatant phase), have been mixed with 180 $\mu$L of supernatant phase (without urea substrate). The Movie fps (frame per second) have been modified to convert the video timescale to 1 minute per second. Scale bar 100 $\mu$m.
- **Supplementary Movie 4. Droplets without urease do not show migration.** In the movie, 20 $\mu$L of droplets phase without enzyme (diluted 1:10, 2 $\mu$L of droplets phase + 18 $\mu$L of supernatant phase), have been mixed with 180 $\mu$L of supernatant phase containing 115 mM urea (final concentration). The Movie fps have been modified to convert the video timescale to 1 minute per second. Scale bar 100 $\mu$m.
- **Supplementary Movie 5. LDH-containing droplets do not display any migration.** In the movie, 20 $\mu$L of diluted droplets phase (diluted 1:10, 2 $\mu$L of droplets phase + 18 $\mu$L of supernatant phase) containing 3.3 $\mu$M LDH have been mixed with 180 $\mu$L of supernatant phase containing 115 mM Urea and 1 mM pyruvate (final concentrations). The Movie fps have been modified to convert the video timescale to 1 minute per second. Scale bar 100 $\mu$m.
- **Supplementary Movie 6. Glucose oxidase-containing droplets do not display any migration.** In the movie, 20 $\mu$L of of diluted droplets phase (diluted 1:10, 2 $\mu$L of droplets phase + 18 $\mu$L of supernatant phase) containing 33 $\mu$M GOx, have been mixed with 180 $\mu$L of supernatant phase containing 115 mM Urea and 50 mM Glucose (final concentrations). The Movie fps have been modified to convert the video timescale to 1 minute per second. Scale bar 100 $\mu$m.
- **Supplementary Movie 7. Migration and merging event between active urease-containing droplets of similar sizes.** In the movie, 20 $\mu$L of of diluted droplets phase (diluted 1:10, 2 $\mu$L of droplets phase + 18 $\mu$L of supernatant phase) containing 1 $\mu$M urease, have been mixed with 180 $\mu$L of supernatant phase containing 115 mM Urea (final concentration). The video screenshots Figure 3B were

obtained from this movie. The Movie fps have been modified to convert the video timescale to 1 minute per second. Scale bar 100 $\mu$m.

- **Supplementary Movie 8. Migration and merging event between active urease-containing droplets of different sizes.** In the movie, 20 $\mu$L of of diluted droplets phase (diluted 1:10, 2 $\mu$L of droplets phase + 18 $\mu$L of supernatant phase) containing 1 $\mu$M urease, have been mixed with 180 $\mu$L of supernatant phase containing 115 mM Urea (final concentration). The video screenshots of Figure 3C were obtained from this movie. The Movie fps have been modified to convert the video timescale to 1 minute per second. Scale bar 100 $\mu$m.
- **Supplementary Movie 9. Migration and multiple merging events between urease-containing droplets driven by pH chemotactical sensing.** In the movie, 20 $\mu$L of diluted droplets phase (diluted 1:10, 2 $\mu$L of droplets phase + 18 $\mu$L of supernatant phase) containing 1 $\mu$M urease, have been mixed with 180 $\mu$L of supernatant phase containing 115 mM Urea (final concentration). The video screenshots of Figure 3D were obtained from this movie. The Movie fps have been modified to convert the video timescale to 1 minute per second. Scale bar 100 $\mu$m.
- **Supplementary Movie 10. Urease-containing droplets show migration and facilitates the formation of a protometabolic simple pathway in a single droplet mediated by the efficient mixing of its content.** 10 $\mu$L of droplets phase (diluted 1:10, 1 $\mu$L of droplets solution + 9 $\mu$L of supernatant phase) containing 1 $\mu$M Urease and 3.3 $\mu$M Alexa fluor 594 labeled-LDH (red) were mixed with 10 $\mu$L of droplets phase (diluted 1:10, 1 $\mu$L of droplets solution + 9 $\mu$L of supernatant phase) containing 1 $\mu$M urease and 3.3 $\mu$M Alexa fluor 488 labeled-PK (green). The droplets mix was imaged during the addiction of 180 $\mu$L of supernatant phase containing 115 mM urea (final concentrations). The video screenshots have been reported in Figure 4A. The Movie fps have been modified to convert the video timescale to 1 minute per second. Scale bar 100 $\mu$m.
- **Supplementary Movie 11. Droplets without Urease show no migration and do not efficiently mix their content.** 10 $\mu$L of droplets phase (diluted 1:10, 1 $\mu$L of droplets solution + 9 $\mu$L of supernatant phase) containing 3.3 $\mu$M Alexa fluor 594 labeled-LDH (red) were mixed with 10 $\mu$L of droplets phase (diluted 1:10, 1 $\mu$L of droplets solution + 9 $\mu$L of supernatant phase) containing 3.3 $\mu$M Alexa fluor 488 labeled-PK (green). The droplets mix was imaged during the addiction of 180 $\mu$L of supernatant phase containing 115 mM urea (final concentrations). The video screenshots have been reported in Figure 4B. The Movie fps have been modified to convert the video timescale to 1 minute per second. Scale bar 100 $\mu$m.
- **Supplementary Movie 12.** Supporting video of the simulation MDMD, Fig. 3B.
- **Supplementary Movie 13.** Supporting video of the simulation SDLD, Fig. 3C.
- **Supplementary Movie 14.** Supporting video of the simulation 4 drops, Fig. 3D.
- **Supplementary Movie 15.** Supporting video of the simulations LDLD, Fig. S10.
- **Supplementary Movie 16. urease-containing droplets with radius $<$40 $\mu$M do not display any migration.** In the movie, 20 $\mu$L of diluted droplets phase (diluted 1:10, 2 $\mu$L of droplets phase + 18 $\mu$L of supernatant phase) containing 3.3 $\mu$M LDH have been mixed with 180 $\mu$L of supernatant phase containing 115 mM

Urea (final concentration). The Movie fps have been modified to convert the video timescale to 1 minute per second. Scale bar 100 $\mu$m.
